# Single-Cell Characterization of Anterior Segment Development: Cell Types, Pathways, and Signals Driving Formation of the Trabecular Meshwork and Schlemm’s Canal

**DOI:** 10.1101/2025.08.15.670438

**Authors:** Revathi Balasubramanian, Nicholas Tolman, Taibo Li, Abdul Hannan, Violet Bupp-Chickering, Karina Polanco, Aakriti Bhandari, Sally Zhou, Marina Simón, John Peregrin, Christa Montgomery, Krishnakumar Kizhatil, Jiang Qian, Simon W.M. John

**Author notes:** Corresponding authors: Revathi Balasubramanian, Simon W.M. John. Authors contributed equally to this work (Revathi Balasubramanian, Nicholas Tolman, Taibo Li).

## Abstract

Morphogenesis of the anterior segment (AS) is crucial for healthy ocular physiology and vision but is only partially understood. The Schlemm’s canal (SC) and trabecular meshwork (TM) are essential drainage tissues within the AS, and their proper development and function are critical for maintaining normal intraocular pressure; abnormalities in either tissue can result in elevated pressure and glaucoma. Here, we use single-cell transcriptomic profiling to provide high-resolution molecular detail of AS development with a particular focus on SC and TM. We report transcriptomes for ∼130,000 single cells at key developmental stages from postnatal day 2 (P2) to P60. We provide the first annotation of cell types across these developmental stages and crucial information about dynamic changes in pathways/gene expression. Further, we trace developmental trajectories for TM cell and SC endothelial cell (SEC) subtypes and determine genes and signaling networks driving their specific cell fates. We demonstrate dynamic changes in signaling interactions between SC and the TM cells during their synchronized development. Collectively, our data lay a deep molecular foundation for AS development that will direct understanding of normal ocular physiology, glaucoma, and other AS conditions.

## INTRODUCTION

Developmental glaucoma (DG) is a group of severe childhood blinding disorders caused by abnormal development of the aqueous humor (AH) drainage structures, including Schlemm’s canal (SC) and the trabecular meshwork^1,2^ (TM). These structures are responsible for regulating AH drainage, and their maldevelopment leads to impaired drainage and elevated intraocular pressure (IOP), which causes glaucoma^3^. DG includes primary congenital glaucoma as well as glaucoma associated with anterior segment dysgenesis (ASD) conditions such as Axenfeld - Rieger syndrome and Peters anomaly^1,4,5^. Thus, the proper development of anterior segment (AS) tissues, including the cornea, conjunctiva, iris, ciliary body, SC, and TM, is essential for vision and ocular health. Genetic studies of patients or animal models with DG and ASD have identified several key genes that are required for proper AS development^5–7^. Notably, variants in genes that cause DG also contribute to adult-onset glaucoma, suggesting that AS developmental pathways have broader relevance to glaucoma pathogenesis^7–9^. Although significant progress has been made in characterizing individual genes, a comprehensive understanding of the molecular regulation of AS development is missing. Major details of its broader developmental trajectory, including highly coordinated processes such as cellular migration, proliferation, differentiation, as well as inter- and intracellular signaling, remain largely undefined. To address this, we provide a global transcriptomic characterization of AS development, while focusing our analyses on the TM and the SC.

The drainage tissues, TM and SC circumscribe the inner surface of the eye wall at the iridocorneal angle, anterior to the iris root. AH first enters the drainage pathway through the TM. In humans, the TM is divided into the uveal, corneoscleral, and juxtacanalicular (JCT) regions. The uveal and corneoscleral regions consist of beams and plates made of various extracellular matrix (ECM) components lined by a monolayer of endothelial-like TM cells. AH primarily flows through intertrabecular spaces that lie between the cell-covered beams. It then passes through the less structured JCT region before reaching the SC. The JCT is comprised of fibroblast-like TM cells embedded in ECM^10^. A similar configuration exists in mice but with less defined TM regions^11^. TM cells regulate IOP through various processes which include cell shape and contractility changes, ECM remodeling, paracrine signaling, limiting oxidative stress, and phagocytosis of cellular debris and other molecules to prevent blockage^12^. Recently using single-cell RNA sequencing, we identified three mouse TM cell subtypes: two beam-like (TM2 and TM3 cells) and one JCT-like (TM1 cells)^13–15^.

Like some other AS structures, TM cells are derived from the periocular mesenchyme (POM). In mice, migration of POM cells into the developing TM begins at embryonic day 10.5 and is complete by postnatal day 6.5. TM cells then undergo differentiation between postnatal days 6.5 and 10. Immature trabecular beams are formed by P10, with more mature beams evident by P14. TM morphogenesis is largely complete by P21, with minor remodeling continuing to around P42^11^. SC and TM development occur synchronously, with each expected to influence the other’s development^16,17^.

SC is a specialized endothelial-lined vessel. It has characteristics of both blood and lymphatic vessels, making it a hybrid endothelial structure^18–21^. SC has distinct inner wall (IW) and outer wall (OW) cells, with different morphologies and transcriptomic states. After exiting the TM, AH enters the SC by passing both between and through the IW endothelial cells, which are specialized for regulating AH drainage. IW cells are elongated and have tight and adherens junction molecules modulating outflow^22,23^. They are enriched for proteins involved in mechano-transduction that have been shown to regulate fluid permeability. The permeability of IW cell junctions is regulated to modulate paracellular fluid flow. Additionally, AH flow pushes IW cells away from their basement membrane attachment points, leading to the formation of extracellular structures called giant vacuoles. This process thins and stretches IW cells, allowing pore formation, where local transient fusion of the basal and apical cell membranes occurs^24^. These transmembrane pores are generally accepted to allow AH to pass through IW cells and into the lumen of SC^25^. From SC, AH drains into the venous circulation via collector channels. These channels originate from the outer wall and connect the canal’s lumen to an intrascleral venous plexus and downstream episcleral veins, ensuring efficient AH outflow (drainage)^24,26^.

SC is derived from limbal veins and develops by a process known as canalogenesis, which combines features of angiogenesis, vasculogenesis, and lymphangiogenesis. In mice, this process starts in early postnatal life. During early SC development (P2.5-P4.5), tip cell (TC) formation, sprouting, and cellular interactions give rise to a rudimentary SC (rSC) structure. In the mid-developmental phase (P6.5-P10), the nascent SC expands through endothelial proliferation. Regarding the late stage (P14-P21), differentiation of the IW and OW SECs as well as lumen formation has occurred (by P14), with all major development and remodeling being complete by P21^18–20^.

TM-derived signals are essential for guiding SC development, maintaining SC health/integrity, and regulating IOP. For example, key signaling pathways that control development, maturation, and function of SC [e.g. VEGFA/KDR (VEGFR2), VEGFC/FLT4 (VEGFR3), ANGPT1/TEK, SVEP1/TEK] utilize ligands produced by TM cells^16,18,20,27^. Importantly, mutations in TEK and other signaling pathway genes contribute to both developmental and adult-onset glaucoma^9,28^, highlighting the importance of studying ocular development to understand the molecular basis of diseases that occur throughout life.

Although recent studies have provided information about individual genes involved in TM and SC development, the molecular mechanisms that drive their development are largely not understood. Single-cell technologies are advancing the understanding of TM and SC biology^13,14,29^, including our recent characterization of three distinct subtypes of TM cells in mice^15^ and the transcriptomes of IW and OW SECs^21^. Despite these advances, many aspects remain unclear, including: 1) the temporal order of TM and SEC subtype development, 2) the molecular events driving TM subtype development; 3) the molecular events driving the hybrid identity of SECs; 4) the nature and interdependence of TM and SC signaling interactions during iridocorneal angle development, 5) the nature of paracrine signaling from other cells types such as immune cells in TM and SC development, and 6) how these mechanisms can be harnessed for therapeutic strategies in glaucoma. A comprehensive understanding of transcriptomic dynamics in the various cell types of the developing iridocorneal angle over the course of TM and SC development is urgently needed.

In this study, we performed single-cell RNA sequencing on the developing mouse AS to understand key developmental processes and the order of events with a major focus on TM and SC. We define for the first time dynamic gene expression changes across the development of both TM and SC. These changes reveal new signaling pathways and molecular regulators, such as YAP/TAZ and Apelin signaling, that are active during SC formation and maturation. We identify the developmental trajectories of TM subtypes and provide a predictive model for their development. At the molecular level, we find that TM and SC development are closely synchronized. Additionally, we identify a novel set of trophic and signaling factors produced by TM cells and other AS cell types that influence SC during its formation and maturation. Together, these findings provide a wealth of new molecular details and lay the groundwork for developing future glaucoma therapies.

## RESULTS

### A single cell atlas of the developing mouse anterior segment

To understand the molecular ontogeny and coordination of SC and TM development, we performed single-cell RNA sequencing of dissected mouse limbal strips (limbal tissue enriched for AH outflow tissues) at key ages of development (Figure 1a, b). We generated ∼130,000 single-cell RNA sequencing profiles of the AS with an average of ∼18,500 cells sequenced at each age. In total across all ages, we profiled 5,846 endothelial cells, of which 1,417 were SECs and 64,683 periocular mesenchyme/neural crest-derived cells, of which 22,799 were TM cells. Combining across all ages, we identified seven broad classes of cells (periocular mesenchyme/neural crest derived cells, epithelial cells, ciliary body (CB) /iris/ retinal pigment epithelial cells (RPE), endothelial cells, immune cells, progenitors, and neurons) (Figure 1c, d, Figure S1). Here, we focus primarily on the development of the TM and SC cell subtypes, providing only an overview of the other developing AS cell types to establish context. Due to the complexity in thoroughly addressing all cell involved in AS development, they will be discussed in detail in subsequent manuscripts.

**Figure 1:**
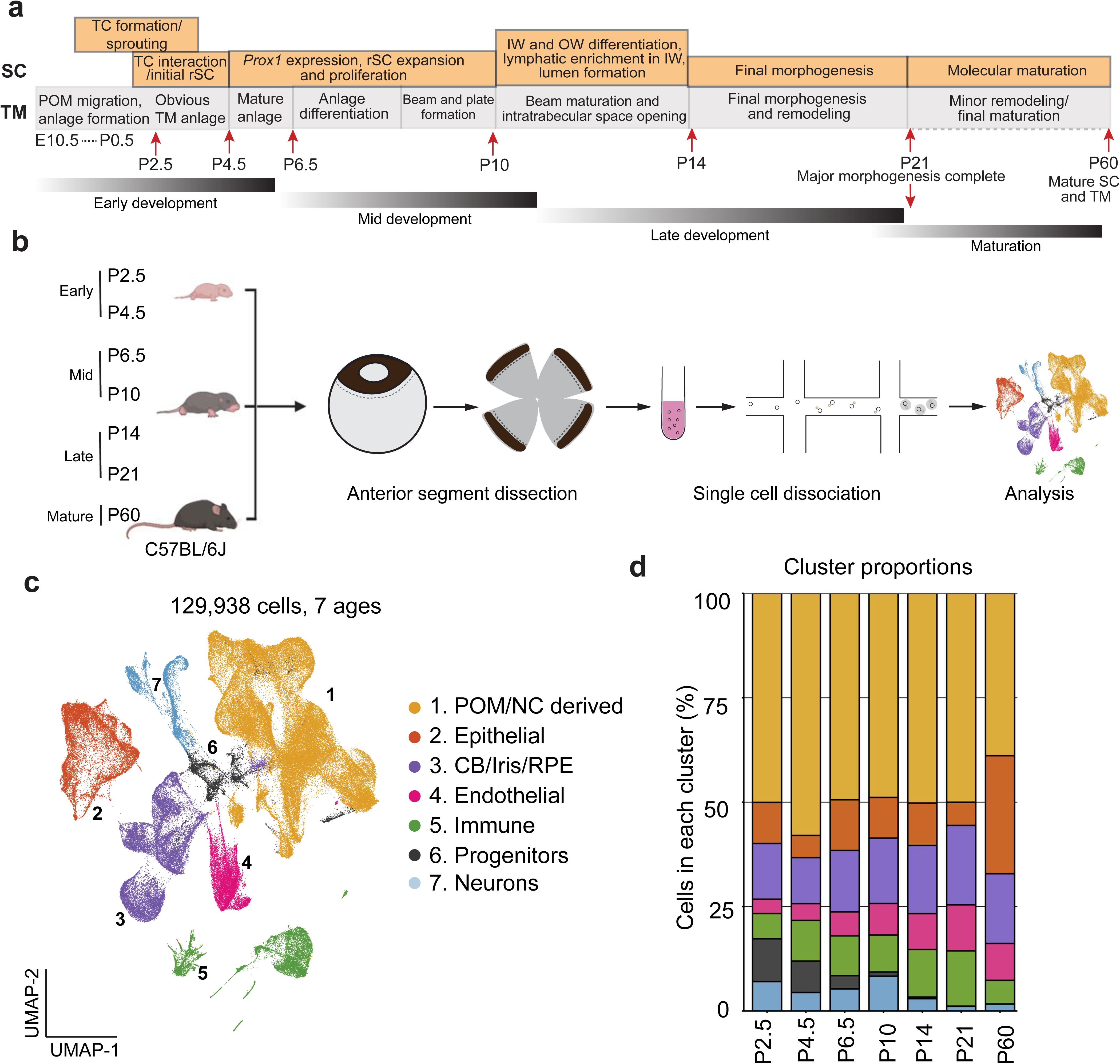
Overview of the single-cell RNA-sequencing experiments. **a)** Diagram of developmental milestones of SC and TM cells in mice. Red arrows indicate timepoints of sample collection. **b)** Diagram of single-cell RNA-sequencing experimental design. Dotted lines indicate cutting planes when dissecting the anterior segment and limbal regions. **c)** Uniform Manifold Approximation and Projection (UMAP) representation of the single cell transcriptomes from all timepoints colored by annotated cell types. **d)** Barplot showing the proportion of each cell type identified across the sampled developmental stages. Some caution is needed in relating to exact *in vivo* proportions due to isolation and processing procedures that can alter cell type proportions.

### Periocular mesenchyme/neural crest-derived cell types

The TM is derived from the periocular mesenchyme/neural crest cells - cluster 1 (C1) while SC is derived from endothelial cells - cluster 4 (C4) (Figure 1c, Figure S1). The periocular mesenchyme (POM)/neural crest (NC)-derived cluster of cells (C1) is characterized by the signature genes *Tfap2b* and *Pitx2* (Figure S2). This cluster consists of five major developing cell clusters (Dev1–Dev5), three previously defined^15^ TM cell subtypes: TM1 (*Chil1^high^, Myoc^high^*), TM2 (*Pgf^high^, Crym^high^*), TM3 (*Acta2*/α-SMA^high^*, Lmx1b^high^*), and other POM/NC-derived AS cell types: Schwann cells (*Egfl8^high^*), pericytes (*Rgs5^high^*), ciliary muscle (*Myh11^high^*), keratocytes, sclera (*Cd34^high^*), iris stromal cells (*Edn3^high^*), and corneal endothelial cells (*Car3^high^*) (Figure 2a - c, Figure S2). The developing clusters are largely present between P2.5 and P10 (Figure 2b, d)^30–33^ along with AS cell types such as Schwann cells, pericytes, ciliary muscle, keratocytes and scleral cells that have already differentiated by P2.5^34,35^ (Figure 2d, e).

**Figure 2:**
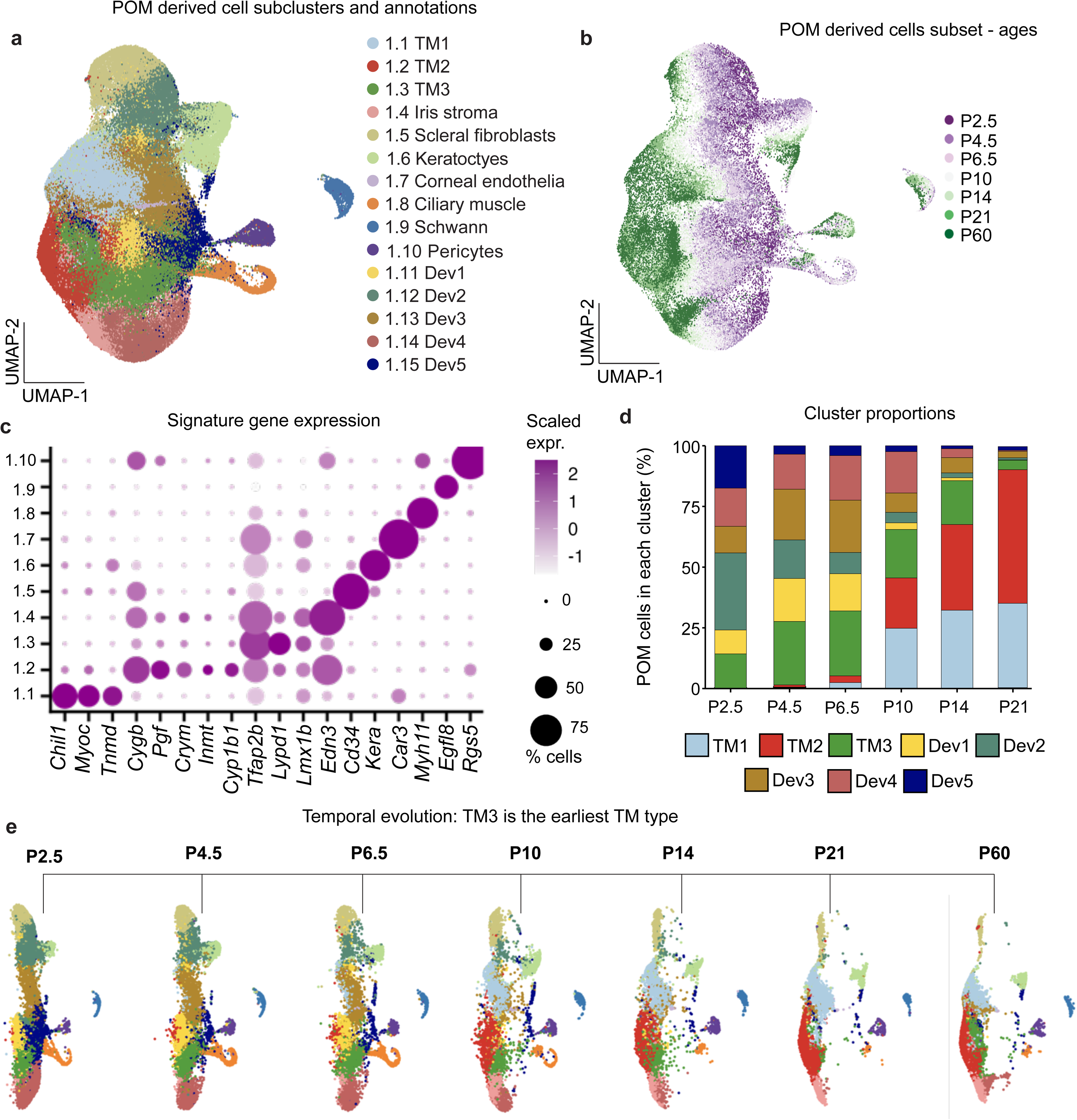
POM/NC-derived cell type analysis. **a)** UMAP representation of POM-derived cells colored by high-resolution cell-type annotations. **b)** UMAP representation of POM-derived cells colored by sample timepoint. **c)** Dotplot showing the expression of key POM marker genes (columns) in each cell type, where size of dots indicates fraction of genes expressed and color indicates average level of expression. **d)** Barplot showing the fraction of each POM/NC- derived cell type across timepoints until P21.Data were integrated across ages, P60 omitted as integration with developing clusters obscured mature cell types at this age. Un-identified technical variation resulted in fewer TM3 being profiled at P21. **e)** UMAP representation of POM/NC derived cells at each timepoint, colored by their cell type annotations (same as in a).

### Development and differentiation of the TM and its cellular subtypes

Morphological changes: We have previously characterized the sequence and timing of morphological changes during TM development using tissue sectioning^11^. Here, we re-evaluated TM development using anterior segment whole-mounts (Figure S3). This allows visualization of the developing TM in 3D around the entire eye. This analysis confirmed our previous conclusions with no changes, hence only a few example images are shown. Immediately after POM migration is complete and the TM anlage is fully formed (by P6.5), the anlage appears as a thick layer of densely packed cells with variable nuclear morphology (rounded and elongated) that are not highly ordered. As differentiation proceeds, nuclei become elongated and begin to become organized along developing beams (P10). By P14, inter-trabecular spaces are clearly evident with beams and spaces becoming more defined by P21 (Figure 3a-c, Figure S4).

**Figure 3:**
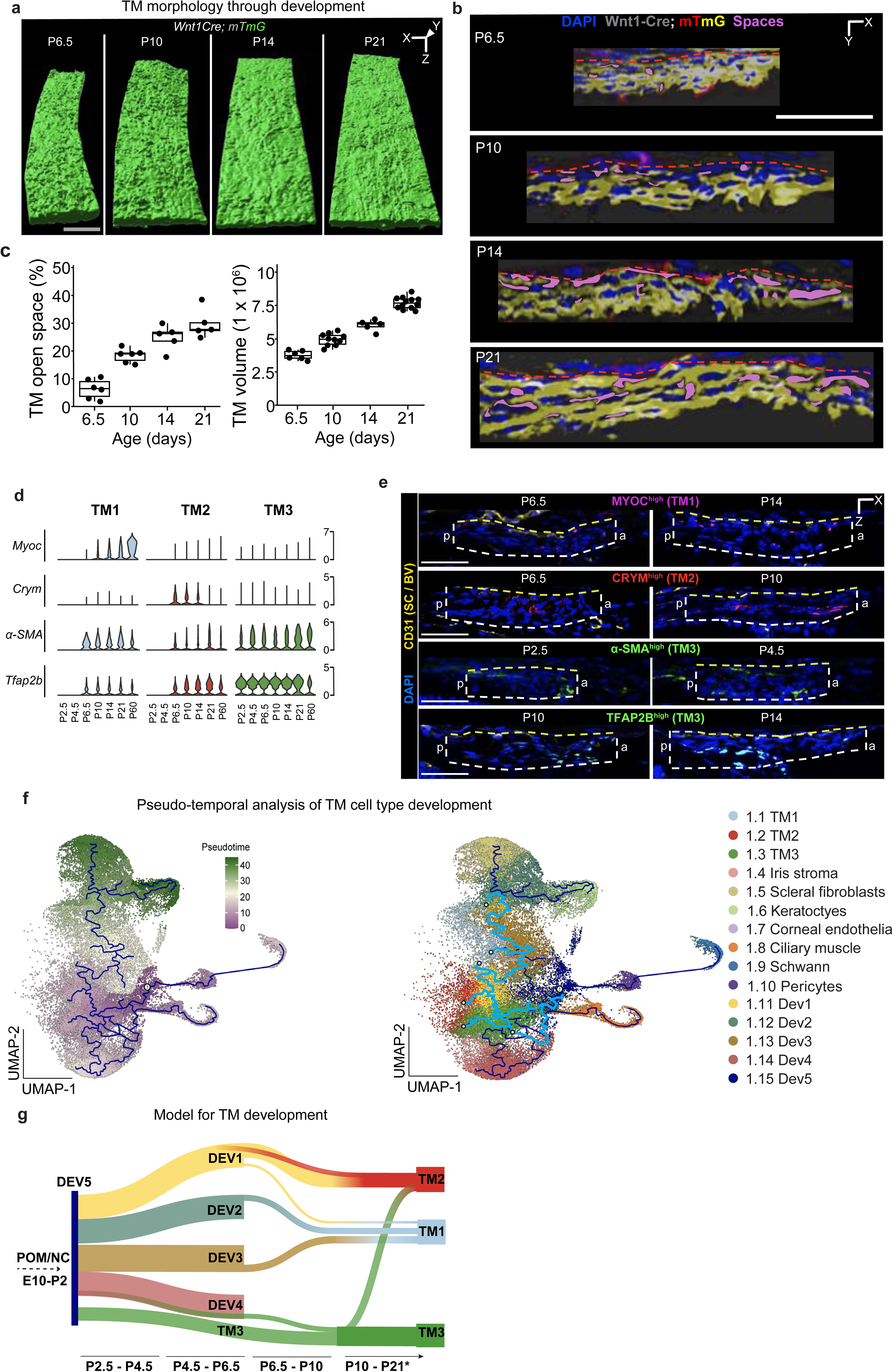
Analysis of TM cell development. **a)** 3D view of a segment of the TM across ages represented by the Surface mode in Imaris. The morphology of the SC facing side of the TM is shown. It becomes progressively more ordered along the longitudinal Y axis of the structure as development proceeds. With obvious beam like striations at P21. Scale bar = 50μm **b)** Visualization of a 1 um thick optically sectioned portion of the TM from a 3D TM reconstruction in the orthogonal XZ plane. Nuclei (DAPI, blue), cell borders (GFP, yellow), and open spaces (magenta, pseudocoloring) of the TM are shown across different developmental stages. At P6.5, the TM is dense with closely packed cells and nuclei (both rounded and elongated). Minimal spaces are evident between the cell borders. As differentiation proceeds (by P10) and inter-trabecular beams and spaces form (P14), the nuclei become elongated and more organized, with increasing volume of space between cells. Clear organization of elongated nuclei and robust inter-trabecular spaces are evident by P21 as major morphogenesis of the TM is complete. Scale bar = 50μm, red dotted line indicates TM border. **c)** Quantification of TM open space and volume with age. **d)** Violin plots showing expression distributions of genes enriched in TM cell subtypes across sample timepoints. **e)** Marker expression across developmental timepoints (immunofluorescence) MYOC – magenta, CRYM – red, a-SMA – green, TFAP2B – green (appears cyan because nuclear co-localization with DAPI). CD31 and yellow dotted lines mark the presumptive SC border. White lines mark TM cells. a: anterior, p: posterior. **f) Left** UMAP embedding of POM/NC-derived cells between P2.5-P10, colored by pseudo-time (Monocle) where the root node was chosen manually as a random cell in Dev5 cluster and labeled in the plot (see main text).**Right:** UMAP representation with monocle pseudo-time trajectory colored by cell types. **g)** Sankey diagram summarizing the development of three TM cell subtypes from clusters Dev1-Dev5.

Differentiation and developmental trajectories: TM3 is the only TM subtype clearly identifiable with robust numbers at P2.5. TM1 and TM2 cells are robustly identified beginning at P10. However, some cells of both subtypes emerge at earlier ages (primarily P4.5-6.5, Figure 2d, e). Because only a very small number of TM1 and TM2 cells are detected at early stages of development (specifically 5 and 65 TM1 cells, and 10 and 97 TM2 cells at P2.5 and P4.5, respectively), the significance of their roles at this time (particularly at P2.5) remains unclear. To further explore this, we sought to correlate the changes in expression of TM subtype-enriched markers as seen in our dataset during development, using immunofluorescence (Figure 3d,e). TM3-enriched markers (α-SMA and *Tfap2b*) were expressed in a subset of TM cells at all examined ages, including P2.5 (α-SMA, Figure 3d). μ-crystallin (*Crym*, enriched in TM2) and *Myoc* expression (enriched in TM1) were detected in TM cells at P6.5, with more robust expression found at P10-P14 (Figure 3d, e). *Myoc* expression was not detected in the developing TM before P6.5 and *Crym* was variably detected at P2.5 and P4.5, possibly because it is expressed in Dev1 cells (Figure S4).

We next analyzed the molecular transformations occurring during TM development. To understand the full array of developmental trajectories for each of the developing clusters of cells leading to TM1-TM3 subtype differentiation, we performed a pseudo-time analysis on cells combining ages from P2.5 (maximum representation of developing clusters) to P10 (all three TM subtypes are first robustly identified). We set a root node in the Dev5 cluster of cells because 1) Dev5 is predominantly present at P2.5 and reduces with each subsequent age (Figure 2a, b, d) and 2) hierarchical cluster tree indicates that Dev5 is closest in identity to Schwann cells and pericytes, which are the earliest cells to differentiate directly from the NC preceding the differentiation of the POM lineage of cells (Figure S5). Starting from the root node, multiple trajectories lead to the differentiation of various TM subtypes as well as other AS cell types. According to pseudo-time trajectory, TM3 is the first defined TM cell subtype, followed by TM2 and lastly TM1 (Figure 3f).

Combining our pseudo-time analysis, cluster tree analysis, and *in vivo* marker analysis, we predict a model for the development of TM subtypes from the Dev1-Dev5 cell clusters (Figure 3g). In this model, the Dev5 cell cluster represents the early NC/POM precursor group that generates the Dev1-Dev4 cell groups. TM3 cells that develop from Dev5, are present from early developmental stages and continue to mature over time. Dev4 also gives rise to new TM3 cells at early developmental ages. Dev2 primarily gives rise to TM1 and other AS cell types starting at early developmental stages, while Dev3 contributes to TM1 formation during mid-developmental stages. Dev1 predominantly contributes to the formation of TM2 cells during early developmental stages but also participates in TM1 development during mid-developmental stages. During mid- and late developmental stages, TM3 also contributes to TM2 development.

Pathways enriched in developing TM cell subtypes: At early developmental stages, TM3 cells are enriched in genes involved in RNA splicing and metabolism, as well as cellular response to vascular endothelial growth factor stimulus. This indicates that, as early as P4.5, TM3 cells are not only differentiating but are also providing signals that are crucial to support SC development (Figure S6). At mid- developmental ages, TM2 and TM3 cells are enriched in genes involved in tissue morphogenesis, sharing features with kidney development and signaling (e.g., integrin binding, fibronectin binding, TGFβ receptor binding) and possibly reflecting common roles in “drainage” of fluids. At late developmental ages, all TM subtypes are largely involved in regulating protein stability, cell-cell signaling, collagen binding, and ECM maintenance indicative of the tissue remodeling at these ages and the maturing functions of TM cells in sensing and responding to changes in IOP (Figure S6).

### Development of other heterogeneous cell types

Sub-clustering the epithelial cell cluster (C2, signature genes *Krt12, Krt5*, Figure S7) characterized a developing epithelial cluster that contained cells predominantly from P2.5 to P6.5. C2 also included corneal and conjunctival epithelial cells, nascent and mature limbal stromal cells, limbal melanocytes, and proliferating epithelial cells^30–33,36,37^. Sub-clustering the third cluster containing cells from the ciliary body and the iris (C3, *Mlana, Tyrp1* signature genes, Figure S8) defined two subtypes of iris stromal cells, as well as distinct populations of iris pigmented epithelium, iris and ciliary body smooth muscle cells, pigmented and non-pigmented cells of the ciliary body, melanocytes, retinal pigment epithelial cells, and a pool of progenitor cells^37–39^. Sub-clustering the immune cells-containing cluster (C5, *Ptprc, Hbb-bs* signature genes, Figure S9) molecularly defined two classes of macrophages, erythrocytes, immune fibroblasts, neutrophils, T cells, B cells, and limbal stem cells^13,32,39,40^. Lastly, sub-clustering progenitor cells^41^ (C6, *Mki67, Top2a*, predominantly from P2.5, P4.5, and P6.5; Figure S10), defined progenitors of retinal neuronal lineage, mesenchymal lineage, epithelial and melanocytic lineage, endothelial lineage, as well as NC cells.

### Endothelial cells during limbal development

The endothelial cell cluster (C4, markers *Egfl7* and *Cdh5* Figure S11), sub-clustered into five major cell types: vascular or blood endothelial cells (BECs - *Cxcl12^high^*, *Slco1a4^high^*)^42^, vascular progenitors (VPs, *Aplnr^high^*)^43^ and collector channel endothelial cells (CECs – *Ackr1^high^*), Schlemm’s canal endothelial cells (SECs – *Selp^high^*, *Npnt^high^*)^21^, proliferating endothelial cells (PECs – *Top2a^high^*, *Mki67^high^*) and lymphatic endothelial cells (LECs – *Lyve1^high^*, *Pdpn^high^*) (Figures 4a, b)^14,21^, with different subsets being predominant at specific ages (Figure 4c, d). As expected, VPs and PECs were largely present at the early and mid- developmental ages. Mature BECs are present at all ages. Mature SECs were present starting at mid- but mostly at the late and mature developmental ages (Figure 4c, d). An unbiased clustering approach did not separate the VPs and CECs into distinct clusters. This may result from: (1) the transcriptome of CECs retaining a predominantly vascular identity that closely resembles that of VPs, preventing them from crossing the resolution threshold for distinct clustering; and (2) limited statistical power to resolve CECs as a separate cluster due to their relatively low abundance compared to other endothelial cell types. LECs were identified across all ages, which is unsurprising because LECs are generated and established by early developmental ages.

**Figure 4:**
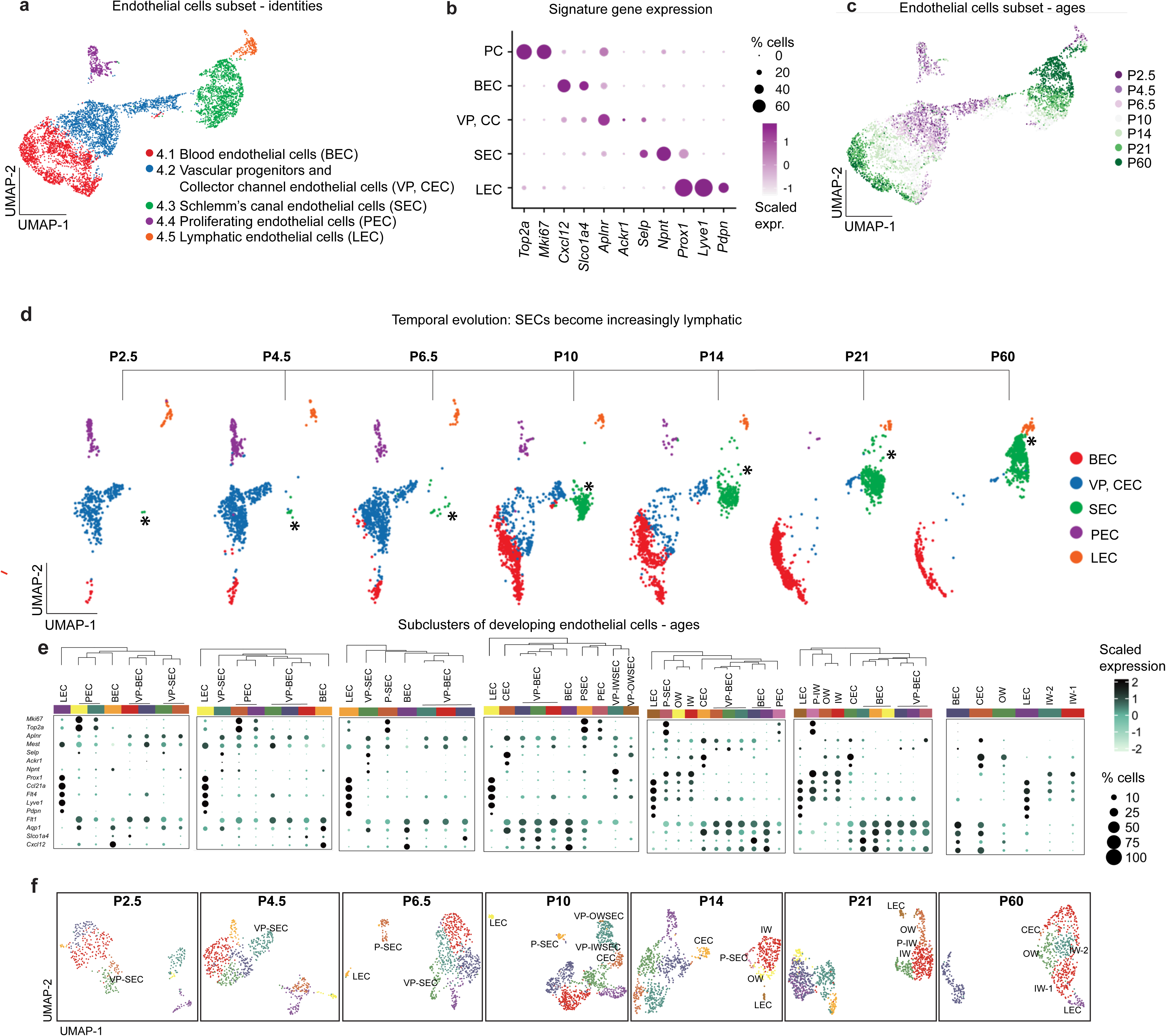
Analysis of SEC development. **a)** UMAP representation of endothelial cells colored by cell subtypes. **b)** Dotplot summarizing expression of marker genes across endothelial cell types where dot size indicates fraction of cells expressing each gene and color indicates average level of expression. **c)** UMAP representation of endothelial cells colored by sample timepoints. **d)** UMAP representation of endothelial cells colored by high-resolution cell type annotations across individual sample timepoints. Asterisk indicates the emergence of Schlemm’s canal endothelial cell cluster (SECs). **e)** Independent analysis of endothelial cells across developmental timepoints – dotplots of expression of key marker genes across each timepoint for each of the dynamically changing endothelial cell types. **f)** Dimensional reduction (UMAP visualization) and cell type annotation as defined separately for each timepoint.

### Differentiation and molecular maturation of SECs

Morphological changes: Previous studies have described the morphological changes and sequence of events involved in the development of the SC. Here, we found no differences in the developmental events compared to what has already been reported using whole mounts^20^ and so we did not use figure space to reiterate these events.

Molecular changes: A closer look at SECs revealed changes in both their number and their UMAP location with age. Unbiased clustering initially placed developing SECs close to the vascular progenitors on the UMAP space at early and mid-developmental ages (see asterisk P4.5, Figure 4d). There is a gradual increase in the number of developing SECs as they proliferate, differentiate, and mature with age (Figure 4d). As development proceeds, SECs become molecularly more similar to LECs and so gradually move closer to them in UMAP position, eventually being immediately adjacent to LECs at P60 (see asterisks P21, P60, Figure 4d). UMAP projections compress high-dimensional transcriptomic data into two dimensions, inevitably distorting the true geometry of the data manifold. Distances and proximities in UMAP space might not reliably reflect the actual transcriptional similarity or developmental distance between cell types. That said, biologically meaningful trends can still emerge despite such distortions, though their interpretation depends heavily on prior knowledge of expected cell type relationships, requiring ground truth labels to judge whether a given embedding is informative. In our case, the spatial relationship between UMAP clusters is in agreement with our previous findings that SECs originate from the blood vessels but acquire a lymphatic-dominant hybrid identity as they mature^20^. Concurrently with SEC development, there is a decline in the VP population. By P21, remaining cells in VP cluster, are primarily CECs as evident by the increase in the expression levels and percent of cells expressing *Ackr1* and *Selp*, markers for CECs (Figure S11). Proliferating ECs (PECs, C4.4) are still present in significant numbers at P14 and persist at P21, but at reduced numbers (Figure 4d). PECs still enter G2M/S phase at P21 and so undergo cell division well into the late stages of AS development (Figure S11). PECs were not detected at P60. This supports a longer developmental time window for ECs in the AS than was previously known.

### Development of inner wall and outer wall SECs

When studying the development of SEC subtypes, we performed clustering separately for each developmental timepoint rather than integrating all datasets across ages prior to clustering. This decision was based on three key considerations: (1) global integration across timepoints can lead to underrepresentation or misclassification of rare or transient cell types that are present at only specific developmental stages; (2) many integration methods prioritize shared features across datasets, potentially obscuring genes with age-specific expression patterns and limiting downstream differential expression analysis within individual clusters; and (3) integration is particularly challenging in the context of dynamic developmental trajectories, where cells may not align cleanly across timepoints due to ongoing changes in cell state.

Vascular progenitors are situated at the UMAP “junction” between presumptive BECs and SECs (Figure 4a, 4d) and contribute to the development of both BECs and SECs. VPs express *Aplnr* and *Mest* at higher levels compared to other ECs. The VPs driving BEC fate (VP-BEC) express vascular genes such as *Flt1*, *Aqp1*, *Slco1a4* and *Cxcl12* ^42,44^ between P2.5 and P14. The VPs driving SEC fate (VP-SECs) lowly express *Prox1* and *Selp* (a marker for OW SECs) at P2.5 (Figure 4e). VP-SECs express *Npnt (*a marker *for* IW SECs) at P4.5, indicating that the developing SECs are becoming further differentiated. Proliferating ECs express both *Npnt* and *Selp* at these early ages (proliferation markers *Mki67* and *Top2a*). VP-SECs are readily detected at P6.5 but are rare or absent at P10, while proliferating SECs (PSECs) are still detected at P10. The most common SEC subtypes at P10 are IW and OW cells that have down-regulated VP (*Aplnr*) and PEC (*Mki67*, *Top2a*) markers. PSECs are still present at P14, while at P21 proliferation genes are primarily expressed in *Npnt*-expressing proliferating IW cells (P-IW). As proliferating OW cells were not detected, this suggests that OW SECs largely generate, differentiate, and mature before P21, while IW SECs continue to be generated at P21. IW cells continue to undergo molecular maturation after P21 with many changes by P60 (Figure 4d-f). At P60, there are well-differentiated clusters of BECs, CECs, LECs, and SECs (*Npnt* ^high^ *Ccl21a* ^low^ IW-1, *Npnt* ^low^ *Ccl21a* ^high^ IW-2, and OW) (Figure 4f).

From our data at P60, it is clear that while all SECs have a mixture of BEC/LEC molecular characteristics (and they are all dominantly lymphatic, including expression of lymphatic master genes *Prox1*^45,46^ and *Flt4*^47^) there is a gradient of cell state between lymphatic and blood vascular EC identity. Accordingly, SC has lymphatic-dominant IW SECs and moderately-lymphatic-dominant OW SECs (Figure 4f, S11). CECs have a more blood vascular biased identity than OW SECs possibly owing to retention of vein-like characteristics during origination (Figure S11). CECs intimately associate with/merge into the outer wall at ostia where collector channels open and emerge from SC.

### Trajectory analysis and immunohistochemistry temporally order the emergence of IW and OW SECs

To understand the development of SEC subtypes from vascular progenitors, we ordered the endothelial cell cluster programmatically according to pseudo-time (Figure 5a). Our unbiased trajectory analysis followed by immunohistochemistry demonstrates the emergence of nascent OW SECs before nascent IW SECs.

**Figure 5:**
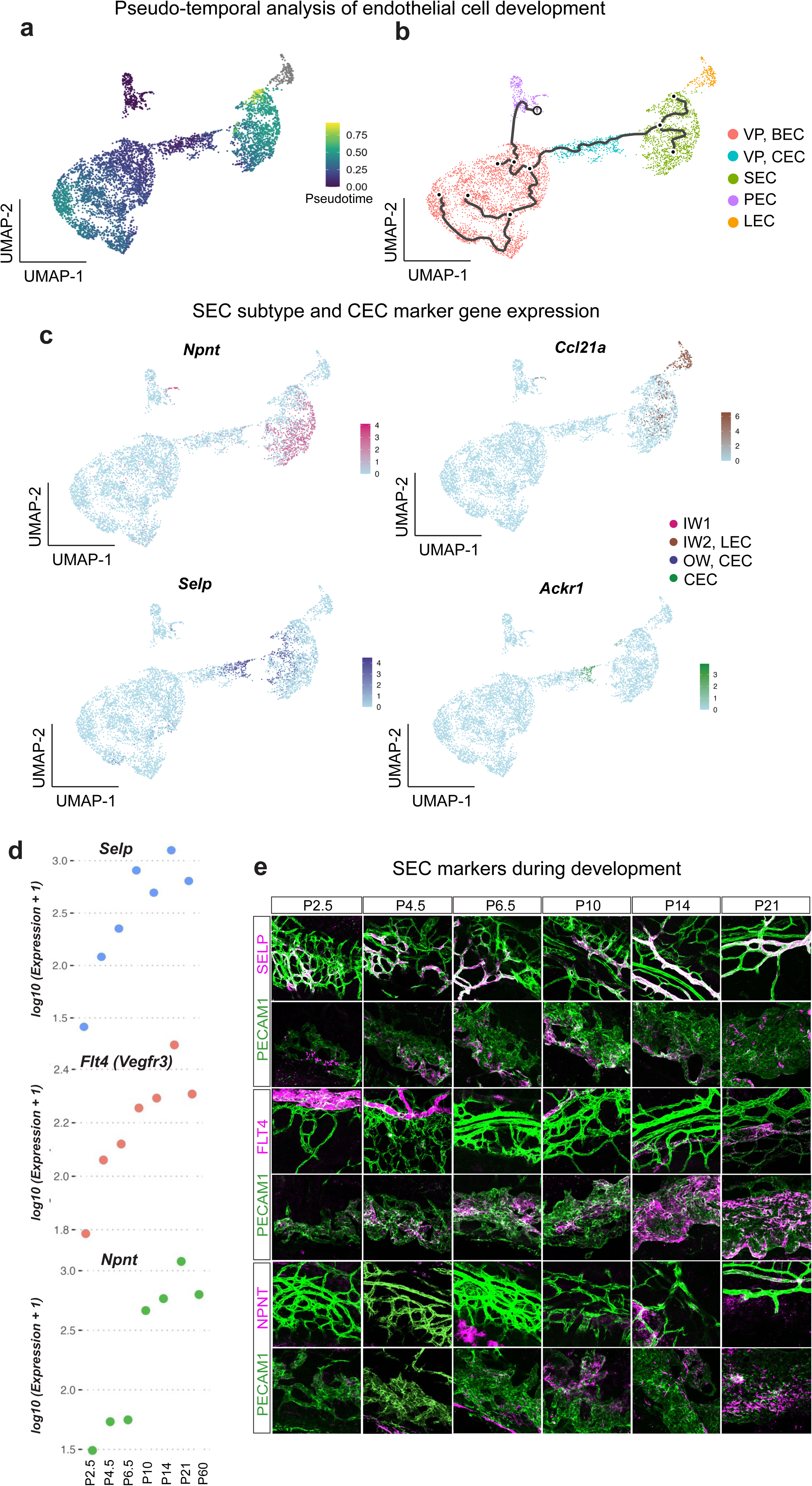
Differentiation of IW and OW of SECs. **a)** UMAP representation of endothelial cells colored by developmental pseudotime inferred by Monocle. **b)** Monocle trajectory superimposed on the UMAP representation of endothelial cells colored by cell type annotation, root node being programmatically determined. **c)** Visualization of marker gene expression plotted on UMAP embedding of endothelial cells. **d)** Scatter plot showing the average expression level of marker genes in a combined population of VP and SECs across sample collection timepoints. **e)** Immunofluorescence labeling of key SEC marker genes across developmental and adult timepoints, top panels showing the vascular plane, bottom panels showing the SC plane from the same Z-stack.

The PEC cluster contained cells with the maximum number of early cells (P2.5 – P6.5) and was programmatically determined to have a pseudo-time of 0.0. A root node, indicating the least differentiated cell state, was assigned at random within this cluster. PEC and VP cells within the BEC and CEC clusters contain the bulk of cells with an earlier pseudo-time, while BEC and SEC clusters contain the bulk of cells with a later pseudo-time (approaching 1.0). The LEC cluster was not reachable by the root node (as LECs are present at all ages) and so was excluded from pseudo-time by the program (assigned an infinite pseudo-time) (Figure 5a). There were multiple branch points from the root node that likely correspond to cell fate switches (Figure 5b). The trajectory of VPs had two major branches: 1) a major node in VP, BECs with three end points likely differentiating into arteries, capillaries, and veins (differentiated BECs), and 2) three end points in SECs likely - OW/CECs, IW1, and IW2 SECs (Figure 5b, c). *Ackr1* (CEC) and *Selp* (CEC and OW) expressing cells are at an earlier pseudotime compared to *Ccl21a* (IW2, LEC) and *Npnt* (IW1) expressing cells (Figure 5c, d, Figure S11). We looked at the overall expression levels with chronological time in VPs and SECs only. *Selp* (CEC and OW marker) expression is higher at earlier chronological time as compared to *Npnt* (IW marker) and *Flt4* (higher and robust expression in IW) (Figure 5d pseudo-time, 5e chronological time). Using immunohistochemistry, we also found that *Selp*-expressing CEC and OW SECs differentiate first from VPs followed by *Npnt* and *Flt4*^high^ IW cells (Figure 5f). *Selp* expression is present in some cells of both the limbal vascular bed and the proliferating chain of rudimentary SC cells in the interstitial zone as early as P2.5. *Npnt* and *Flt4* expression starts at P6.5 and is obvious by P10 (Figure 5f). Overall, this demonstrates that the greater the lymphatic bias of a cell type, the later it emerges (CEC, most blood vascular first; OW, intermediate next; then IW SECs, most lymphatic last).

### Gene expression changes underlying early SC development

To characterize the gene regulatory networks underlying the various stages of SC development, we first analyzed the biological processes enriched in VPs and SCs at early developmental ages (Figure 6a, b). In VPs, there was enrichment in pathways related to endothelial cell development, migration, and proliferation, among others (Figure 6a). In early SECs, as expected, biological processes such as histone modification and regulation of cellular macromolecule biosynthesis (needed for cell division and growth) were enriched (Figure 6b, complete list in S12, S13). In further exploring genes in the enriched endothelial cell proliferation and endothelium development processes in VPs, we found that genes involved in endothelial sprouting and migration^44,48,49^ (such as *Aplnr*, *Flt1*, *Kdr*, *Nrp1*, *Clec14a*, and *Jmjd6)* were high in VPs during early development and subsequently reduced with time (Figure 6c). In contrast, these genes started to be elevated at around P4.5 in developing SECs (when rudimentary SC cells are sprouting, proliferating, and migrating to form chain of SECs) and reaching peak expression at P10. This timing suggests their continued roles in regulating SC development, maintenance and function as supported by the well-known vaso-regulatory roles of *Flt4* and *Kdr* (both VEGF receptors, Figure 6d)^50^. Notably, *Aplnr* (a key gene involved in tip cell development and sprouting angiogenesis in various other vascular systems^51,52^) is at its highest expression in developing SECs at P10 (mature SEC differentiation first evident). *Aplnr* expression begins to be reduced after this age as SECs continue to be differentiated and decreases more substantially after P21 (Figure 6d). This implies crucial roles for *Aplnr* at all stages of SC development.

**Figure 6:**
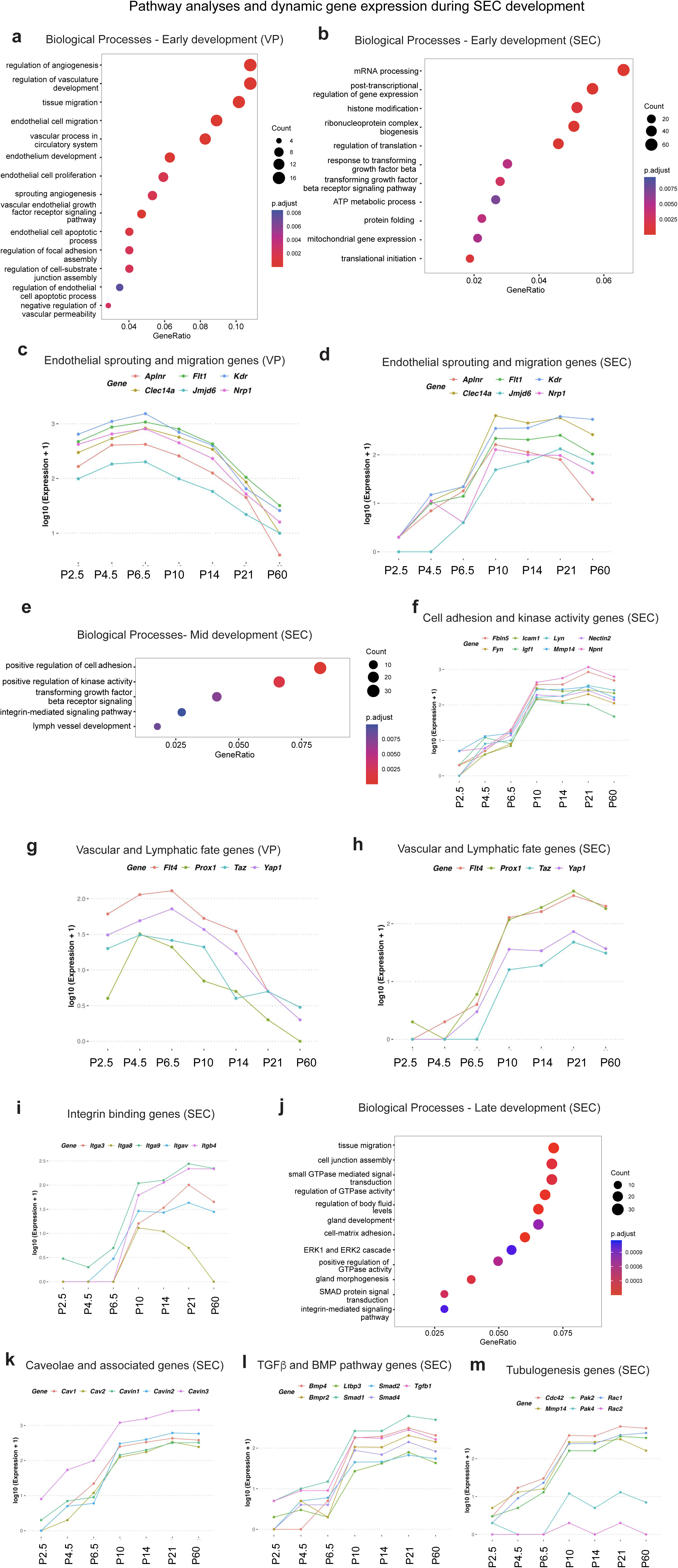
Dynamic gene expression analysis during Schlemm’s canal endothelial cell development. **a-b)** Gene ontology (GO) enrichment of biological process genes selectively expressed during early development in a: VPs; b: SECs. In all GO plots, x-axis represents the fraction of genes in each pathway which are also identified in the corresponding differentially expressed gene set. Colors indicate adjusted p-value of enrichment, and dot size indicates number of genes in the pathway. **c,d)** Average expression level of genes in selected GO biological processes enriched in VPs and SECs during early development, across sample collection timepoints. **e)** GO enrichment of genes expressed during mid-developmental timepoints in SECs. **f-i)** Average expression level of genes in select processes enriched in VPs or SECs across sample collection timepoints**. j)** GO enrichment of genes expressed in SECs at late development. **k-m)** Average expression level of genes in select processes enriched in SECs across sample collection timepoints.

### Gene expression changes underlying the transition from VPs to lymphatic-biased SECs

SECs originate from limbal veins and transition to a mixed blood vascular and lymphatic phenotype that is biased towards a lymphatic identity. To understand the gene regulatory activities underlying the blood vascular-lymphatic transition, we looked closely at genes involved in blood vascular and lymphatic development in VPs and SECs. At mid-developmental ages, pathways regulating cell adhesion, kinase activity, integrin-mediated signaling, and lymph vessel development are enriched in SECs (Figure 6e). In VPs, genes encoding proteins that promote angiogenic blood vessel formation, such as Yes-associated protein 1 (YAP) and WW domain-containing transcription regulator 1 (WWTR1/TAZ)^53–55^, are at high levels during early developmental time points but decline with age (Figure 6g). At the same time, the key lymphangiogenic regulators *Prox1* and *Flt4* are also elevated in VPs during early stages, peaking around P4.5 and P6.5 when rudimentary SC cells are seen. In developing SECs, *Prox1* and *Flt4* are already upregulated by P6.5 but undergo their steepest expression rise between P6.5 and P10 - a critical window during which proliferating SECs acquire a lymphatic identity (Figure 6h). Although *Yap1* and *Taz* expression also increase over this interval, their upregulation is less pronounced than that of *Prox1* and *Flt4*. Importantly, all four genes maintain their high but differential expression into adulthood, suggesting that the dynamic interplay between these pro-lymphatic and pro-blood vascular programs drives the mixed vascular yet lymphatic-biased identity of SECs.

### Maturation and adult function of SECs

Our data provide information on many other genes and pathways active in SECs during SC development and maturation to the adult state. Mentioning a few: 1) Genes important for cell-cell adhesion and cell-matrix adhesion such as *Fbln5*, *Mmp14*, and *Npnt* ^56–59^(Figure 6f). 2) Genes that mediate immune to endothelial cell interactions like *Icam1*, *Igf1*, *Fyn*, *Lyn*, and *Nectin2* ^60–63^(Figure 6f) - with a marked increase in their expression at P10. 3) Integrins and other mechano-sensors that are expected to be important in developmental signaling and in regulating aqueous humor drainage though junctions^64,65^. Several integrin family members are also elevated by P10 with the expression of most integrins trending up with age. *Itga8* is an exception in becoming down-regulated with developmental progression (Figure 6i). The key SEC protein NPNT binds to ITGA8 to mediate developmental signaling in other tissues, but during SC development *Itga8* is downregulated as *Npnt* expression rises, making their interaction unlikely. 4) Caveolae: Caveolae with their component proteins and modulators are important in mechano-transduction-based signaling^66–68^. The *Cav1* and *Cav2* genes, which are associated with elevated IOP and POAG^69,70^, are robustly expressed as early as P6.5 with near peak expression levels by P10. *Cavin3* has the highest level of expression compared to other caveolar genes and has significant expression as early as P2.5 (Figure 6k). 5) Metabolism and mitochondria: In agreement with our previous study^21^, mature SECs are enriched in biological processes and molecular functions related to metabolic and mitochondrial function. Molecular functions pathway enrichment in mature SECs includes proton and active ion transmembrane transporter activities, electron transfer activity, cytochrome-c oxidase activity and biological processes such as aerobic respiration, oxidative phosphorylation, ATP metabolic process, and mitochondrial respiratory chain complex assembly (Figure S12, S13). This suggests that maintenance of SC and regulation of aqueous humor outflow by SECs are energy intensive processes. 6) TGFβ)/BMP/SMAD signaling. Transforming growth factor beta signaling – highly implicated in primary open angle glaucoma (POAG)^9,71,72^ – is enriched in SECs throughout development (Figure 6b, 6e, 6j). Various important genes in this signaling (*Tgfb1, Ltbp3, Smad1, Smad2,* and *Smad4*) increase by P10 and are at their highest expression levels from P14 and onwards, with *Bmp4* and *Bmpr2* mirroring this expression pattern (Figure 6l). BMPs are previously implicated in SC and TM development and IOP regulation^73–77^.

### Gene expression changes underlying lumen formation

The cells forming the early SC structure form a continuous chain of cells around the ocular limbus well before the lumen of SC is formed^20^. Schlemm’s canal begins to form a lumen with inner and outer walls at P10, with the lumen only evident at some local positions^11,20^. Between P10 and P14, the lumen forms around the entire canal with SECs in the IW and OW gaining their distinct mature morphologies. The molecular mechanisms underlying lumen formation or tubulogenesis in SC are not previously characterized. Our grouped data for SECs from P14 to P21 reveals substantial expression of genes and pathways known to mediate endothelial cell tubulogenesis and lumen formation in other systems ^78,79^. Upregulated tubulogenesis pathways within SECs include the regulation of GTPase activity, ERK1 and ERK2 cascade regulation, vascular development regulation, and integrin-mediated signaling (Figure 6j). Looking at specific genes in these pathways, we found that tubulogenesis genes and regulators, including *Cdc42*, *Mmp14* (encoding MT1-MMP), *Pak2,* and *Rac1,* ^78,79^ were up-regulated between P10 and P14, the major period when lumen formation occurs. Although *Pak2* and *Rac1* increase by P10, expression of their paralogs *Pak4* and *Rac2* remains low, implying minimal involvement (Figure 6m). At the same time, we start to see expression of pericyte-marker genes including *Rgs5, Kcnj8, Vtn* (vitronectin) and *Des* (desmin, Figure S14)^80^. It is possible that pericytes were recruited but were not identified as a distinct cell cluster because their numbers are relatively small compared to other cell types. Pericyte recruitment is a known feature of tubulogenesis in other vessels^78^. Further experiments are needed to assess the role of pericytes in SC development.

### Developmental signaling from TM to SC

It has long been assumed that the development of SC is guided by signaling from the TM, with recent studies defining initial signaling pathways. To provide a more global analysis, and the role of signaling from specific TM cell subtypes, we used our transcriptomic data to evaluate signaling at different stages of SC development. Predicted ligand-target interactions suggest that TM3 cells are a major driver of SC development at early developmental stages, expressing *Vegfa*, *Vegfc*, and *Angpt1-* key factors previously shown to be important in SC development by functional tests. Several other genes for soluble factors such as *Tgfb1*, *Tgfb2*, *Tgfb3*, and *Edn3* are also expressed by TM3 cells during early stages of SC development. Our predictive interaction analysis also identifies other growth factors such as *Fgf18*, *Fgf21*, *Igf1*, and *Hbegf* that are made by TM3 cells and have defined molecular targets on SECs (Figure 7a, b, Figure S15 provides ligand-receptor pairs). As the SC and TM continue their synchronized development, and aligned with TM cell subtype differentiation, developing TM1 and TM2 cells assume larger roles in the release of growth factors such as *Vegfa* and *Angpt1* by mid developmental stages and have largely taken over from TM3 by P10 (Figure 7c, d). With further developmental progression, TM1 and TM2 cells produce most of the growth and maintenance factors (Figure 7e, f). *Vegfc*, a critical growth factor in SC development (receptor *VegfR3*/*Flt4*), is dynamically produced by all three TM cells between P6.5 to P21 and beyond (Figure S16 contains expression levels of all ligands across TM subtypes), consistent with a critical role in the development and maintenance of SC.

**Figure 7:**
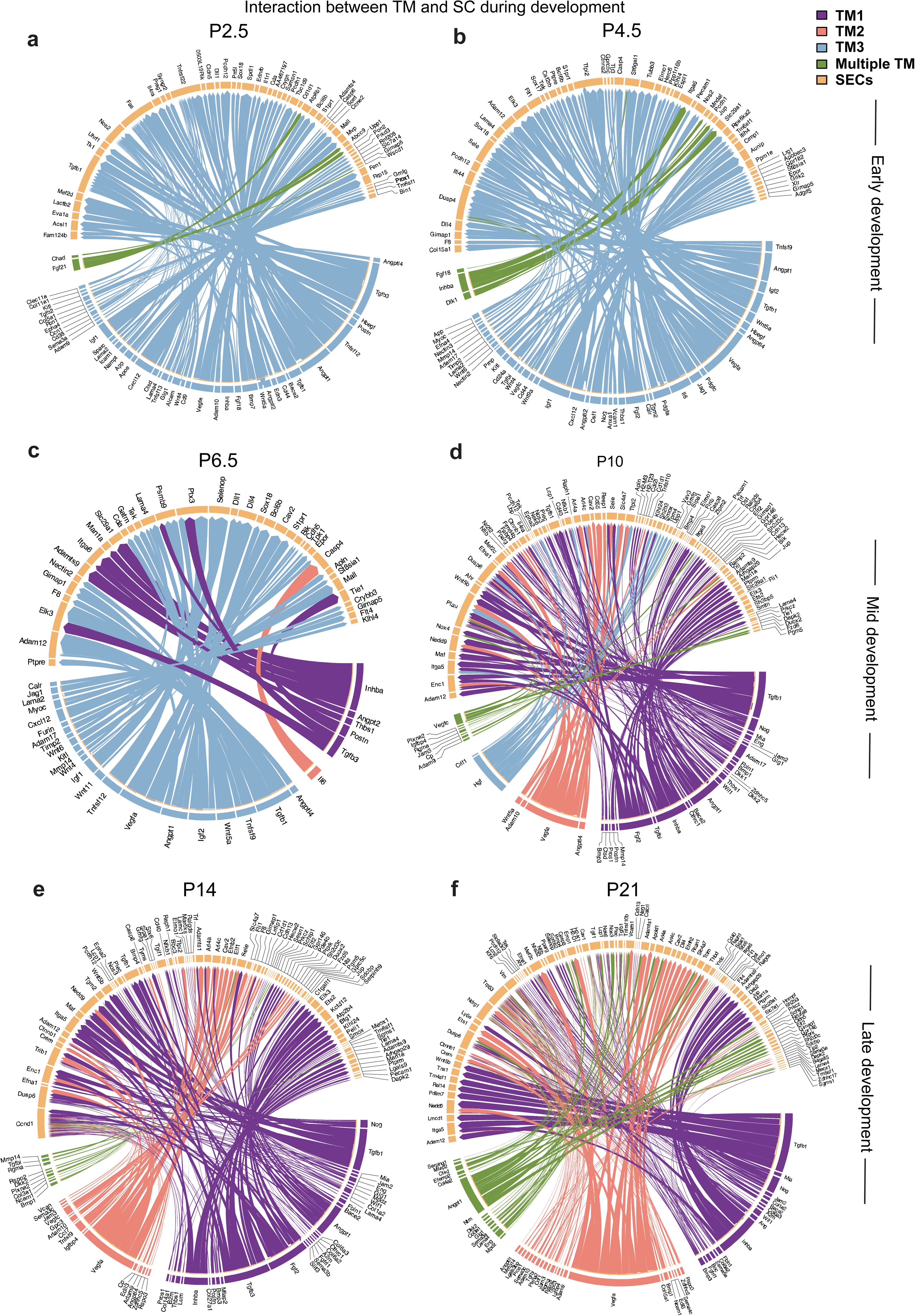
Ligand-target interaction analysis between TM and SC cells. **a-f)** Circos plots demonstrating inferred interactions between TM (sender) and SE (receiver) cells across sample collection timepoints stratified by early (P2.5 and P4.5), mid (P6.5 and P10), and late (P14 and P21) timepoints.

### Signaling between other cell types and SC/TM

We also determined predicted ligand-target interactions between other cell types that are likely to modulate SC or TM development and were not included above (Figures S17, S17 and S19). Additional cell types were macrophages (known chaperoning and signaling roles in SC development, likely important in TM development and modulate outflow)^64^, vascular progenitors, pericytes (reside on collector channels and although they are not shown to modulate SC development, they may have a local role), and SECs (may signal to other SECs and to developing TM). A comprehensive representation of these signaling relationships between all cell types and at all key ages is beyond this paper and so select developmental ages are shown. Key signaling pathways with targets on developing and mature SECs include *Bmp4*, *Jag1*, *Dll4* and *Tgfb1* (ligands produced by VPs, Figure S17) as well as *Ackr3, Bmpr2,* and *Kdr* (ligands produced by macrophages, Figure S18). Macrophage-produced ligands with receptors on TM cells include *Il1r1, Itgav*, and *Bmpr2* (Figure S19). Importantly, macrophage modulators/chemo-attractants are produced by developing SECs (e.g., *Csf1*, *Csf2*, *Csf2rb*, *Csf3b Cntf, Cxcl12)* (Figure S17) indicating that developing SECs themselves may influence their own development and function by modulating the role of macrophages (Figure S18). The developing and mature SECs can also potentially signal to adjacent TM cells. This pattern, where most SEC signaling at P10 and P21 appear to target TM1 cells (Figure S19), is consistent with the TM1 subtype being positioned directly adjacent to the inner wall of SECs^15^. Overall, our datasets and our ligand : target analyses provide the most comprehensive information on local intercellular signaling during TM and SC development to date.

### Genes associated with POAG and elevated IOP during SC and TM development

Lastly, we looked at genes associated with elevated IOP and POAG in GWAS studies^69,71,72,81–83^ (Figure S20). The 5 developing clusters (Dev1-Dev5) have a similar enrichment of GWAS genes as mature TM cells, supporting possible developmental, adult, or both developmental and adult roles for these glaucoma genes. Pathway analyses of genes implicated in patients with developmental glaucomas show an enrichment in pathways involved in SC and TM development. SC pathways include the VEGF signaling pathway, receptor tyrosine kinase binding pathway, and the lymph vessel morphogenesis pathway. TM pathways include neural crest development, mesenchymal cell development, and collagen fibril organization. A unifying theme was that the expression levels of these genes neared peak values at early to mid-developmental stages (as early as P4.5 in TM and P10 in SC) and maintained similar levels during late development and after maturation (Figure S20). This is consistent with the involvement of some genes in both developmental and adult-onset glaucoma.

## Discussion

### Deep molecular characterization of TM and SC development remains needed

In 2001, Smith et al.^11^ published a comprehensive assessment of mouse iridocorneal angle morphogenesis. They outlined the series of cellular and morphological transformations. Most detail was provided on TM development, from the formation of the trabecular anlage by POM-derived cells to final TM maturation. They resolved a controversy in the field by showing that programmed cell death was not a major remodeling process during angle development. At the time, SEC markers were not known, limiting characterization of early SC development. A major step in understanding SC development was in three closely spaced reports in 2014^18–20^. These studies included the first detailed description of the entire process, from initial tip cell sprouting from limbal veins to mature SC. Kizhatil et al.^20^ used the highest-resolution confocal imaging and distinguished IW from OW SECs. They named the entire process of SC development canalogenesis, laying out the then novel developmental sequence, which sequentially combined features of angiogenesis, lymphangiogenesis, and vasculogenesis. These three studies identified the mixed vascular and lymphatic features of SECs driven by *Prox1* expression^18–20^.

Comparing the IW and OW, Kizhatil et al.^20^ also discovered that the lymphatic molecules are more enriched in the IW which mediates AH drainage and predicted the importance of TEK (alias TIE2) signaling in SC development and maintenance based on its high expression level. Key regulatory molecules were also identified (including VEGFA/VEGFR2 alias KDR), VEGFC/VEGFR3 alias FLT4). These important studies stimulated the field with subsequent studies expanding the players in TM and SC development (e.g., *Tie1*, *Angpt1, Svep1, Vtn, Bmp4, Foxc2, Jag1,* etc. and implicating some in developmental glaucomas)^16,27,64,73,84–86^. Despite these and other important advances, a comprehensive molecular characterization of TM and SC development remains lacking. Critical molecular details have been needed regarding the differentiation, maintenance, and identity of distinct cell types and subtypes involved in TM and SC formation and function.

### The first comprehensive molecular characterization of AS development

Here, we provide the first comprehensive evaluation of TM and SC development using single-cell resolution transcriptomics. We identify and molecularly characterize the participating cell types at all key developmental stages providing a novel atlas that captures dynamic cellular changes throughout the developmental ontogeny of TM and SC. Although we focus on TM and SC in this paper, our datasets capture the key expression changes in all major cell types of the developing AS over the same time period when other AS tissues also develop and mature. Our molecular data emphasize the degree of coordination between SC and TM development with close alignment of key milestones in cell proliferation, differentiation, and remodeling. We highlight key signaling molecules produced by various TM subtypes, and other cell types that may induce and coordinate developmental changes in SC as well as regulate and maintain SC function. Critically, the molecules and pathways we identify - along with their temporal dynamics align closely with the known developmental sequence of SC and make strong biological sense in the context of the processes underway. Importantly, our findings provide an unprecedented wealth of molecular information that will advance future understanding of both SC and AS development as well as support the creation of novel therapies for glaucoma.

### Developmental trajectories, cell types, and switches in trophic factor production guiding TM and SC development

#### TM cells

Previously TM cell differentiation was thought to occur from P6 onwards^11^. Our data clearly show that TM3 cells are present in robust numbers by P2.5, when the trabecular anlage is still forming and before any morphologically obvious differentiation or remodeling of the anlage. This is also a stage where initial sprouting of tip cells from limbal vessels is occurring as SC development commences. Thus, cues from the developing SC are unlikely to induce TM3 differentiation but TM3 cells may be involved in induction of SC.

Based on our pseudotime, trajectory, and real chronological time analyses, we propose a new model for the development and differentiation of TM cell subtypes. Our model and data reflect an initial development of TM3 cells, followed by TM2 cells, then lastly TM1 cells. This model shows complex lineage relationships with some TM subtypes having contributions from different developmental precursor cells at different stages of development. A recent study used single-cell RNA sequencing to profile developing POM cells in rats at late embryonic and early postnatal stages^41^. Integrating their dataset with ours across species will provide a more comprehensive understanding of TM development beginning with early POM. While these new details are valuable and begin to inform developmental as well as directed differentiation strategies for TM (for treatment purposes), our datasets will allow detailed follow up analyses including integrating more POM-derived cell types (including the iris stroma, ciliary muscle, keratocytes, sclera) to more fully understand how various cells types communicate and influence each other’s development.

#### SECs

In agreement with previous studies^18–20^, our data demonstrate SECs have a mixed vascular phenotype with characteristics of both blood and lymphatic vessels. We show that they develop from more blood vascular precursors (not surprising as they arise from veins) and then gain their lymphatic bias as development progresses (based on our pseudotime, trajectory and real chronological time analyses). The relatively more blood vascular outer wall cells are identified first (nascent OW with marker gene expression evident at P2.5), with IW cells that have strongest lymphatic bias differentiating last (nascent IW with marker gene expression evident starting at P6.5).

Our data advance understanding of vascular signaling ligands in SC development. Previous studies by Aspelund et al.^18^ and Kizhatil et al.^20^ showed the importance of *VegfR2 (Kdr)* and the *Vegfc-VegfR3 (Flt4)* signaling axis for SC development. Here, we document the production of VEGF and other ligands by the TM cell subtypes with strong shifts in production between TM subtypes as development proceeds. For example, *Vegfa* is initially expressed by TM3 cells but as development proceeds, and TM1, TM2 cells differentiate, *Vegfa* becomes largely expressed by TM2 cells. *Vegfa* is also expressed by SECs cells themselves. Similarly, *Angpt1* production, a ligand that signals through the TEK receptor and modulates SC development and function, shifts from TM3 cells in early development to largely TM1cells at later stages. *Angpt1* is also expressed by the developing SC cells (in agreement with Thomson et al.^27^). Thus, the expression of trophic factors that are necessary for SC development is dynamic and sequential at both the level of cell-intrinsic expression by SECs, and cell-extrinsic expression by specific subtypes of TM cells.

### Molecules driving the mixed vascular, lymphatic-biased fate of SECs

SC is one of six known hybrid vessels that exhibit both blood and lymphatic endothelial characteristics. The others include the liver sinusoids, high endothelial venules of secondary lymphoid organs, penile cavernous sinusoids, the ascending vasa recta in the kidney, and the remodeled spiral arteries of the placenta^87,88^. Despite their unique dual identity, the precise molecular cues that govern this hybrid fate determination remain largely unknown. Our analyses determine that during early stages of SC development, VPs exhibit dynamic gene expression patterns that shift from angiogenic towards lymphangiogenic pathways, a shift that is clear in all of our pseudotime, trajectory, and chronological time analyses. During this shift there is upregulation of *Prox1* and *Flt4*, key drivers of lymphatic identity, while expression of *Yap1* and *Taz*, key drivers of angiogenic identity, is retained. These lymphatic drivers become more highly expressed than the blood vascular drivers, suggesting that the balance of these factors determines the prevailing mixed vascular but lymphatic biased identity of SECs. However, further experiments are needed to directly test this. This balance may be one of several regulatory systems influencing the hybrid fate of SECs. Future studies combining lineage tracing with epistasis analyses will be crucial in understanding the precise mechanisms controlling this process.

### Key signaling and intercellular interactions among AS cell types

During development, it has been shown that heterogeneous groups of cells communicate and influence each other’s developmental programs. ASD usually presents with malformations in multiple cell types, underscoring the importance of not studying the development of single cell types in isolation. Here, our data support numerous inter-cellular signaling interactions with potential to influence SC and TM development. A few of the interactions that we identify between TM and SC are: i) EDN3 (ligand) produced by the TM acting upon EDNRB (receptor) expressed by SECs during early development of SECs – altered endothelin signaling has been implicated in various forms of glaucoma^89,90^, ii) *Tgfbi* produced by the TM targeting E-Selectin on SECs during development – variants in *TGFBI* were recently identified in patients with congenital and juvenile onset glaucoma^91^, iii) Periostin (POSTN) released from the TM and directly or indirectly targeting downstream molecules such as TIE1 on SECs – molecules shown to have roles in TM and SC development respectively^84,92^, iv) Insulin-like growth factor 1 (*Igf1*) – a pro-angiogenic factor^93^ released from multiple sources (TM, macrophages, and pericytes) to influence SC development, and v) Multiple BMP ligands from various sources all acting upon SC – BMP2 from macrophages, BMP3 from TM, and BMP4 from SC. A more in-depth investigation of these predicted pathways through functional and molecular studies in mouse models will be essential to test many of the predicted interactions, both informing developmental mechanisms and treatment strategies

### A role for macrophages in SC and TM development

Previous work by Kizhatil et al.^20^ demonstrated that macrophages interact with rudimentary SC cells during early development and likely have a role in chaperoning cellular interactions. This work also demonstrated the close association of significant numbers of macrophages with adult SC. Subsequently, GWAS have highlighted a significant role for immune cells particularly macrophages in modulating the risk of POAG^71^. More recently, it was shown that macrophages can regulate aqueous humor outflow and intraocular pressure^94^, while laser treatments of the TM of glaucoma patients have been suggested to modulate IOP via macrophage recruitment and signaling in the TM^95^. Building on these findings, we find that the developing TM expresses macrophage-related receptors such as *Il1r1* and *Itgav*, indicating that macrophages may influence the development of both SC and TM. In addition, our analysis suggests that SC cells may actively recruit macrophages through the expression of genes such as *Cxcl12, Cntf*, and various interleukins. Furthermore, SC cells express ligands and receptors, including *Csf1, Csf2, Csf2rb,* and *Csf3r*, which are associated with the formation of a macrophage niche. Collectively, our findings suggest a novel role for SECs in recruiting macrophages and they identify candidate genes through which macrophages may influence TM and SC development, with implications for developmental and adult glaucoma. Supporting the importance of these findings, a recent paper demonstrated that vitronectin on macrophages signals through integrin αvβ3 signaling on SECs, with disruption of this signaling resulting in mal-development of SC^64^.

### IOP and POAG GWAS genes are active during development and in adults

We also provide a comprehensive analysis of GWAS gene expression during postnatal anterior segment development. Genes associated with elevated IOP and POAG, such as *Cav1, Cav2, Fbn1,* and *Foxc1,* reach peak expression by P10 during SC development. Similarly, Tgfβ signaling pathway members are also expressed at near-adult expression levels as early as P10 in both the developing SC and TM. Our overlay of the timing and expression level of GWAS genes on specific cell types in the developing and adult anterior segment provides important information for understanding their role in ocular development and glaucoma.

Anterior segment development is a highly coordinated process in which heterogeneous cell types of distinct embryonic origins. Studying any single tissue in isolation therefore omits critical intercellular interactions indispensable for normal development, and fails to capture the multi-tissue basis of conditions like ASD and DG. Here, we present a comprehensive single-cell transcriptomic atlas of the developing anterior segment, identifying transcriptional trajectories, cell fate determinants, and dynamic signaling interactions that drive SC and TM maturation in the context of the broader anterior segment cellular ecosystem. Beyond SC and TM, this atlas provides a blueprint for future studies incorporating additional anterior segment cell types, and the gene regulatory networks uncovered here represent a rich source of candidate disease mechanisms and therapeutic targets for ASD, developmental glaucoma, and related anterior segment disorders. Given the strong conservation of anterior segment and ocular drainage tissue development between mice and humans, these datasets offer a powerful foundation for human studies and targeted therapy development.

## Methods

### Animal husbandry and ethics statement

All experimental animals for single cell RNA sequencing and TM/SC morphology analyses were C57BL/6J (Stock# 664) inbred mice. Additionally, for analysis of TM morphology, we used a combination of the B6.Cg-E2f1Tg(Wnt1-cre)2Sor/J (Wnt1-Cre)^96^ and B6.129(Cg)-Gt(ROSA)26Sortm4(ACTB-tdTomato,-EGFP)Luo/J (mTmG)^98^ strains. We crossed these alleles together as described in Kizhatil et al. ^20^, which labels the SC and surrounding vasculature with Tdtomato (red) and the TM and a subset of surrounding structures with GFP (green). All mice were treated in accordance with the Association for Research in Vision and Ophthalmology’s statement on the use of animals in ophthalmic research. The Institutional Animal Care and Use Committee of Columbia University approved all experimental protocols. Mice were maintained on PicoLab Rodent Diet 20 (5053, 4.5% fat) and provided with reverse osmosis-filtered water. Mice were housed in cages containing ¼-inch corn cob bedding, covered with polyester filters. The animal facility was maintained at a constant temperature of 22°C with a 14-hour light and 10-hour dark cycle.

### Limbal tissue dissection

Pregnant dams were checked daily between 9:00 AM and 12:00 PM for births. The date of birth was recorded as midnight of the day pups were first observed. Mice were collected at the following timepoints: P2.5, P4.5, P6.5, P10, P14, P21, and P60. For each timepoint, eyes from four animals (eight eyes total) were pooled per sample, and two samples were collected per age. All samples consisted of eight eyes, except for one P21 sample, which contained seven eyes due to exclusion of a severely microphthalmic eye. Sex was balanced at each timepoint. Sex was determined by anogenital distance at P21 and P60, and by genotyping at earlier ages (see below). To standardize early timepoint collections, P2.5 and P6.5 harvests were conducted between 7:00–9:00 AM, while all other timepoints were collected between 9:00 AM–12:00 PM.

Eyes were enucleated into fresh Dulbecco’s Modified Eagle Medium (DMEM; Gibco) and dissected in the same medium. All dissection tools and surfaces were treated with RNaseZap (Ambion). For dissection, the posterior segment was removed by cutting between the limbus and sclera, followed by lens removal. Finally, we made a small circular cut in the anterior cornea, removing the middle 1/3^rd^ of the area (P2.5–P14), and the middle 2/3^rds^ for P21 and P60. This change in dissection likely contributed to the reduced number of TM3 cells observed at P21. TM3 cells are enriched anteriorly (at-least in adult) and so are located closer to the corneal cut during dissection of the P21 eyes (which despite being larger than younger ages are still smaller and more delicate to accurately dissect than at P60) and are therefore more likely to be lost. The iris was carefully removed at P21 and P60. The remaining anterior segment tissue (limbal strip) was minced and prepared for enzymatic digestion.

### Single-cell RNA sequencing

Tissues were digested enzymatically (Papain Dissociation System; Worthington Biochemical Corporation) and single cells were loaded into 10× Chromium Single Cell Chips (detailed protocol in Balasubramanian et al. ^21^). Single-cell libraries were generated according to the manufacturer’s protocol. Libraries were sequenced on the Ilumina Novaseq platform. Sequencing data were demultiplexed and aligned using Cellranger software (10× Genomics). Reads were aligned to the mouse mm10 genome.

### Single-cell RNA sequencing data processing

Single-cell RNA-seq data preprocessing was performed following Seurat v4^97^ pipeline with custom modifications. For each sample, we first removed cells with the number of RNA features lower than 100 and higher than 3000, and with more than 20% mitochondrial reads. Cell doublets were further inferred using Scrublet^99^ with default parameters in each sample, and removed from further analyses. We then performed SCTransform v2^100^ with percent mitochondrial reads as covariates to normalize each dataset. Subsequently, we applied a two-step integration workflow by first integrating samples corresponding to each developmental day (resulting in seven integrated single-cell RNA-seq datasets), and then integrating across all days for the final integrated dataset, using FindIntegrationAnchors and IntegrateData functions in Seurat with 3000 features. Subsequent clustering and 2D projection by UMAP were performed using the default Seurat pipeline with 30 principal components. Leiden clustering was used with resolution 0.4 to derive initial clusters. For SC and TM clusters, we performed sub-clustering to improve resolution of cell type definition. Additionally, cell clusters were manually merged into one of the seven coarse cell types (POM/NC derived, epithelial, CB/Iris/RPE, endothelial, immune, neurons, and progenitors).

For each cell type, we identified marker genes using FindMarkers function in Seurat with a negative binomial generalized linear model, requiring genes to be detected in at least 25% of cells with log fold-change >= 0.25. Marker genes were matched with previously published work to annotate cluster identities.

For developing clusters in the POM/NC cluster of cells, we used a combination of strategies to annotate them: Previous studies have defined the molecular identities of adult limbal cell types, including their marker gene expression profiles^14,15^. Developmental clusters were identified based on the absence of clear signature gene expression corresponding to any adult cell type. Frequently, the gene expression signatures of these developmental clusters resembled hybrids of multiple adult cell types. Moreover, all developmental clusters were abundantly present during early developmental stages (P2–P14) but were sparsely detected at later timepoints.

To estimate temporal dynamics of gene expression, for each cell type, we estimated transcriptome changes by aggregating cells collected in the same day, applied linear models with collection day as the independent variable to assess classify genes up- or down-regulated across development. Pseudobulk gene expression was visualized using scatter or line plots with mean expression per day for each gene, as well as heatmap with k-means clustering to group genes with similar temporal profiles across all genes. In addition, we applied Monocle^101^ to reconstruct developmental trajectories using default parameters, and calculated pseudotime to obtain high-resolution dynamic expression patterns for each gene and estimate developmental transition probabilities between cell types. To further dissect the molecular basis of interaction between cell types, we obtained receptor-ligand interaction data from NicheNet^102^ and applied a modified version of LRLoop^103^ using genes expressed in at least 10% of cells in each cell type to infer feedback loops of receptor-ligand interactions for each developmental timepoint. Enrichr^104^ was used to estimate enriched biological pathways in each cluster based on the resulting list of marker genes. ClusterProfiler^105^ was used for gene ontology analyses. For analysis of GWAS genes, using genome-wide association studies of primary open-angle glaucoma^69,71,72,81–83^, we curated a list of genes associated with the disease. Seurat’s AddModuleScore function was applied to assess glaucoma disease risk for each cell type, using default parameters (24 bins for aggregation with 100 control features per analyzed feature). Developmental glaucoma genes used for pathway analysis were curated from relevant literature^6,106–109^.

For hierarchical clustering, TM-containing cells were assessed based on the similarity of the expression of the top 500 marker genes (ranked by P value) of each individual cluster at each age. This analysis filtered out low-expressing genes (expressed in >10% of cells, with a logFC > 0.25 compared to other cells).

### Genotyping XY chromosomes

The presence of X and Y chromosomes was determined by a previously established genotyping protocol^110^. Sex was determined by this genotyping. Genomic DNA was amplified with the forward primer 5’ GATGATTTGAGTGGAAATGTGAGGTA 3’ and reverse primer 5’ CTTATGTTTATAGGCATGCACCATGTA 3’. Genomic DNA was PCR amplified using the following program: 1) 94°C for 2 minutes, 2) 95°C for 30 seconds, 3) 57°C for 30 seconds, 4) 72°C for 30 seconds, 5) repeat steps 2-4 30 times, 6) 72°C for 5 minutes. 6) 72oC for 5 minutes. 5 μl of sample was run on a 1.5% agarose gel. The X chromosome amplifies a 685 base pair fragment, and the Y chromosome amplifies a 280 base pair fragment.

### Preparing eyes for immunohistochemistry

Enucleated adult eyes of postnatal mice were fixed for 1 hour at 4°C in 4% paraformaldehyde (PFA, Electron Microscopy Science, Hatfield, PA) prepared in phosphate buffered saline (1X PBS, 137 mM NaCl, 10 mM phosphate, 2.7 mM KCl, pH 7.4).

#### Whole mounts

Following fixation, the postnatal eyes were dissected out according to a previously established method^20^. Briefly, the anterior part of the eye was cut just posterior to the limbus, and the iris, lens, ciliary body, and thin strip of retina were carefully removed to obtain the anterior eye-cup. The anterior cup includes the cornea, limbus, and sclera. Four centripetal cuts were made to relax the eye-cup and facilitate eventual mounting onto a slide after immunohistochemistry (method below).

#### Frozen Sectioning

Following fixation, a small punch was made through the optic nerve so that eyes would not shrivel during dehydration. Eyes were dehydrated in a 30% sucrose (Sigma-Aldrich, St. Louis, Missouri) in 1X PBS at 4C until eyes sunk to the bottom of a 2ml glass vial. Once sunken, eyes were embedded in optimal cutting temperature embedding medium (Fisher Scientific, Waltham, MA) and flash frozen on dry ice. Eyes were cryo-sectioned at 10μm thickness in the sagittal plane. Sections were evenly spaced throughout the peripheral and central ocular regions.

### Immunohistochemistry

#### On whole mounts

Anterior eye-cups were washed multiple times with 1× PBS to quench residual fixation. The eye-cups were incubated with 3% bovine serum albumin and 1% Triton X-100 in 1× PBS (blocking buffer) 4°C for 2 h in 2 ml glass vials to block nonspecific binding of antibody and to permeabilize the tissue. The anterior cups were then incubated with primary antibodies of choice in 200 µl blocking buffer for 2 d, with rocking, at 4°C. The anterior cups were then washed three times over a 3-h period with 1× PBS. The primary antibodies were detected with the appropriate species-specific secondary antibody (Alexa Fluor 488, 594, or 647 at 1∶200 dilution, Life Technologies, Grand Island, NY) diluted in blocking buffer, which also had 4′,6-diamidino-2-phenylindole (DAPI, 1:1000) (Thermo Scientific, Waltham, MA) to label nuclei. The immunostained eye-cups were washed four times over a 3hour period in 1× PBS. Eye-cups were then whole-mounted on slides in ProLong™ Diamond Antifade Mountant (Invitrogen, Waltham, MA). For primary antibodies used and concentrations, see antibodies table (Supplemental Table 1).

#### On sections

Sections were washed three times with 1× PBS with 0.3% Triton X-100 for 5 m to quench residual fixation. The sections were incubated with 10% Donkey serum (Sigma-Aldrich, St. Louis, Missouri) and 0.3% Triton X-100 in 1× PBS (blocking buffer) at room temperature for 1 hour to block nonspecific binding of antibody and to permeabilize the tissue. Sections were then incubated with the primary antibodies indicated in 200 µl blocking buffer overnight at 4°C. The sections were then washed three times for 5 minutes with 1× PBS with 0.3% Triton X-100. Primary antibodies were detected with the appropriate species-specific secondary antibody (all Alexa 488, 594, or 647 at 1∶1000 dilution, Life Technologies, Grand Island, NY) diluted in 1× PBS with 0.3% Triton X-100 for 2 hours at room temperature. The secondary antibody solution also included 1:1000 DAPI (Thermo Scientific, Waltham, MA) to label nuclei. The immune-stained sections were washed three times for 5 minutes in 1× PBS with 0.3% Triton X-100. Sections were then mounted using Flouromount (Sigma, St. Louis, MO). For primary antibodies with low target antigen abundance, tyramide signal amplification was used according to previously published protocol^15^. For primary antibodies used and concentrations, see antibodies table (Supplemental Table 1). Greater than 20 sections were examined for at least 1 major TM cell subtype marker (MYOC, CRYM, and α-SMA) at all ages (P2.5, P4.5, P6.5, P10, and P14), except for CRYM which was not examined at P14. In addition to α-SMA, TFAP2B was examined as a TM3-enriched marker at P6.5, P10, and P14 (6-10 sections examined per age).

### Microscopy of drainage structures

Microscopy was performed using an LSM SP8 confocal microscope (Leica) using a 63×1.4 NA glycerol immersion objective and 40× 1.1 NA water immersion objective. The Mark and Find mode was used to automate collection of images in Z stacks (across the depth of the limbus from the limbal vessels down to the trabecular meshwork with Schlemm’s canal in between) at various individual overlapping positions along the limbus.

### Postprocessing of Images

#### 3D rendering

We followed the guidelines detailed in Kizatil et al. and Tolman et al. ^15, 20^ for visualizing SC and TM in 3D. Individual confocal Z stacks (.lsm files) were processed directly using Imaris 9.5 (Oxford Instruments, Carteret, NJ). Multi-position Z stacks were first imported into Imaris and converted into Imaris files (.ims files). The resulting Imaris files were then stitched to generate a comprehensive Z stack encompassing the limbus using ImarisStitcher 9.5. The stitch was exported as an Imaris file and processed appropriately. For sections, images were taken from the Z-plane in focus. The snapshot feature of Imaris was used to generate high resolution (1024×1024 pixels, 300 dpi) images for all figures. 3D rendered images utilized maximum intensity projections generated in “Surpass” mode of Imaris. Images were oriented so that structures of interest were visible.

#### SC and TM segmentation and analysis

To analyze the SC and TM planes, specific regions were projected after eliminating extraneous planes by using the Crop 3D function of Imaris. Cropping was performed within 5µm on either side of the structure of interest (see Kizahtil et al.^20^ for details on SC segmentation and Figure S3 for details on TM segmentation). SC and TM structures were defined based on previously characterized 3D anatomy expression of either mT or mG in the Wnt1Cre; mTmG system and other antibodies that are known to label either SC and TM (SC, PECAM and VE-cadherin; TM, α**-**SMA). However, none of our antibodies is specific to either SC or TM; they only label the structure and a small number of surrounding structures. These surrounding structures can then be eliminated based on morphology, leaving a 3D crop or either SC or TM. The TM was defined based on anatomical landmarks as described previously^15^. Briefly, we defined the TM as the region between the inner wall of Schlemm’s canal (SC, outer TM) and the anterior chamber (inner TM), extending from the anterior edge of the pars plana (posterior TM) to the posterior edge of Descemet’s membrane of the corneal endothelium (anterior TM).

#### TM volume and space analysis

The gross 3D morphology of the TM was analyzed using the Surface feature in Imaris to generate a 3D object based on GFP expression in the cropped TM plane. Individual TM layers were examined by digitally cropping a 1 µm-thick section along the longitudinal axis and visualizing nuclei (DAPI) and cells (GFP). A surface object of the TM was generated, and all voxels outside this surface were set to null. GFP-positive voxels (Wnt1Cre+ GFP; TM cells) located within the TM surface were exported as a histogram of 0–255 intensity values by transferring the data from Imaris to ImageJ. Background was determined by measuring the intensity of GFP signal in the space of the anterior chamber within 1-5 μm of the TM surface). The percentage of empty space was calculated by dividing the number of voxels at or below background levels by the total number of voxels above background within the TM surface. The same quadrants and eyes were analyzed as in the other TM analyses across ages. 6-10 whole eyes were analyzed at each age. Due to a degree of unavoidable tissue distortion and compression during processing, the volume of the intertrabecular space is likely an underestimate, especially at older ages where the tissues are more fragile and more prone to such artifacts.

## Data availability

Raw and processed data can be accessed on GEO (GSE315712) and Single cell portal (SCP3301) respectively.

## Funding sources

This project was supported by the Brightfocus Foundation National Glaucoma Research grant G2021007S (R.B), HHMI, the Precision Medicine Initiative at Columbia University, National Eye Institute (NEI) grant R01EY032507, and the New York Fund for Innovation in Research and Scientific Talent NYFIRST - EMPIRE CU19-2660 (S.W.M.J). Partial support was provided by NEI grants R01EY11721, R01EY018606 (S.W.M.J), R01EY032062 (S.W.M.J, and K.K), The Glaucoma Foundation (SWMJ), and R01EY034493 (J.Q). Support also came from start-up funds from Columbia University (R.B, S.W.M.J), and Ohio State University (K.K). Additional support was provided by Research to Prevent Blindness (RPB) Career Development Award (R.B), The Glaucoma Foundation’s grant-in-aid (R.B), Glaucoma Research Foundation Shaffer research grant (R.B), BrightFocus Foundation grant CG2020004 (to S.W.M.J and K.K), and the Knights Templar Eye Foundation career starter grant (A.H), RPB new chair challenge grant to Ohio State University (K.K), and unrestricted departmental award from RPB to Columbia University (R.B, S.W.M.J). Core grant support also contributed: P30EY019007 (Columbia University) P30EY00176 (Johns Hopkins University). RB is a Chang Burch Scholar. S.W.M.J is the Robert Burch III Professor of Ophthalmic Sciences and was an Investigator with the Howard Hughes Medical Institute (HHMI) during the first years of this project. The content is solely the responsibility of the authors and does not necessarily represent the official views on the National Institutes of Health.

## Author contribution statement

R.B designed and performed experiments, analyzed data, wrote the manuscript, and sought and obtained funding. N.T conceived, designed and performed experiments, analyzed data, wrote and edited the manuscript. T.L designed analyses, analyzed data and edited manuscript. A.H performed experiments and edited manuscript. V.B-C performed experiments. K.P analyzed data and edited manuscript. A.B performed experiments. S.Z performed experiments. M.S edited manuscript. J.P performed experiments. C.M edited manuscript and assisted with laboratory and mouse colony management/maintenance. K.K analyzed data and edited manuscript. J.Q designed analyses, oversaw work at Johns Hopkins and edited manuscript. S.W.M.J conceived and oversaw overall study and designed experiments, analyzed data, wrote and edited manuscript, sought and obtained funding. Notes on joint authorships: This was a major project of the John Lab where R.B and N.T partnered on tissue/data collection, analysis, and validation tests with R.B focusing more on Schlemm’s canal while N.T focused more on trabecular meshwork. They discussed analysis needs with T.L who supported their analysis efforts and provided essential analyses and computational expertise throughout the project under the guidance of J.Q. R.B wrote and finalized manuscript with S.W.M.J after starting her own laboratory, with her lab members contributing to new data analyses and validation tests.

## Supplemental figures – legends

**Figure S1.**
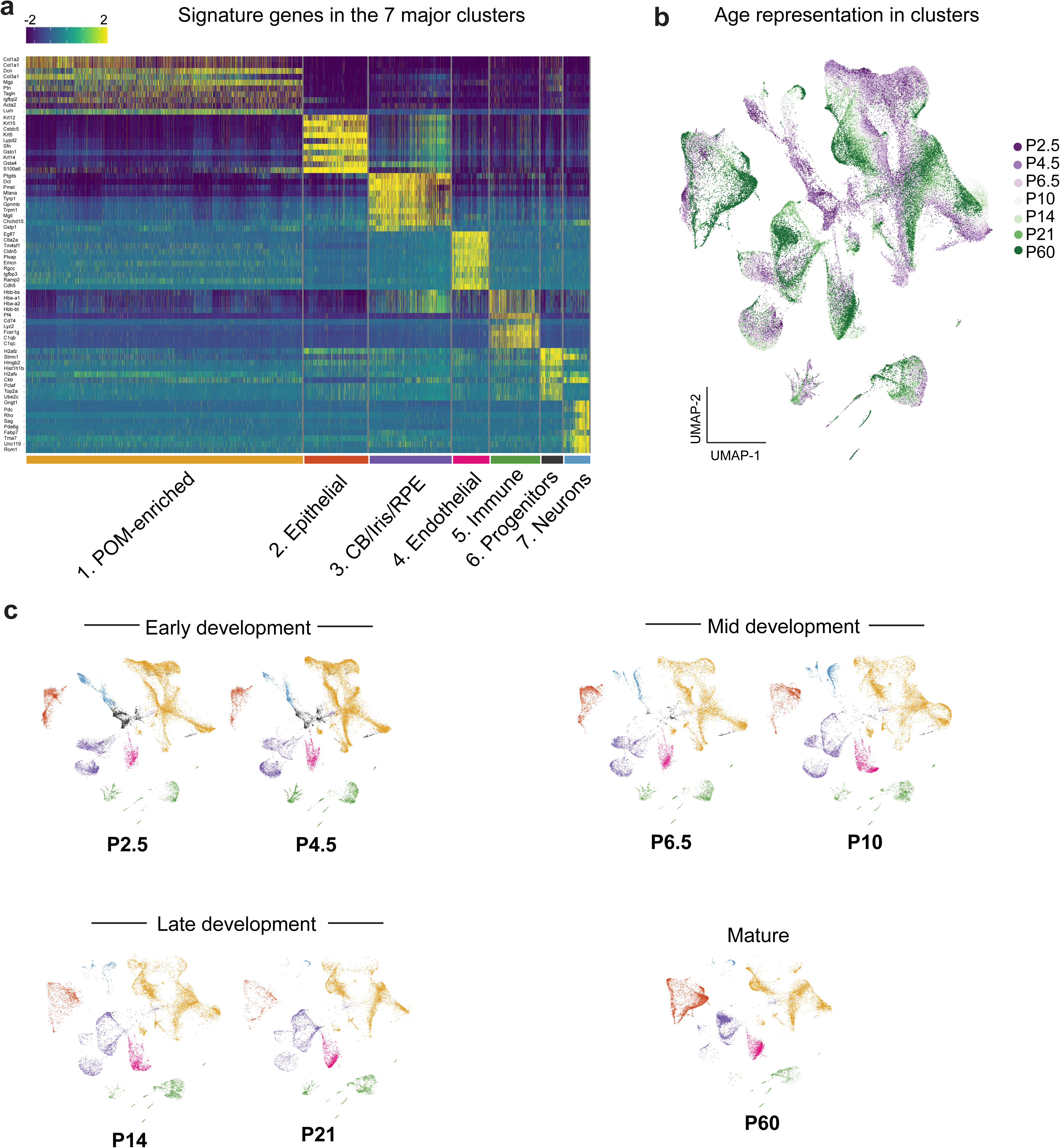
Cell type annotation. **a)** Heatmap showing the expression of signature genes (rows) across single cells (columns), arranged by seven major cell types. **b)** UMAP representation of all cells colored by sample collection timepoints, **c)** UMAP representation of all cells colored by cluster identity at each timepoint.

**Figure S2.**
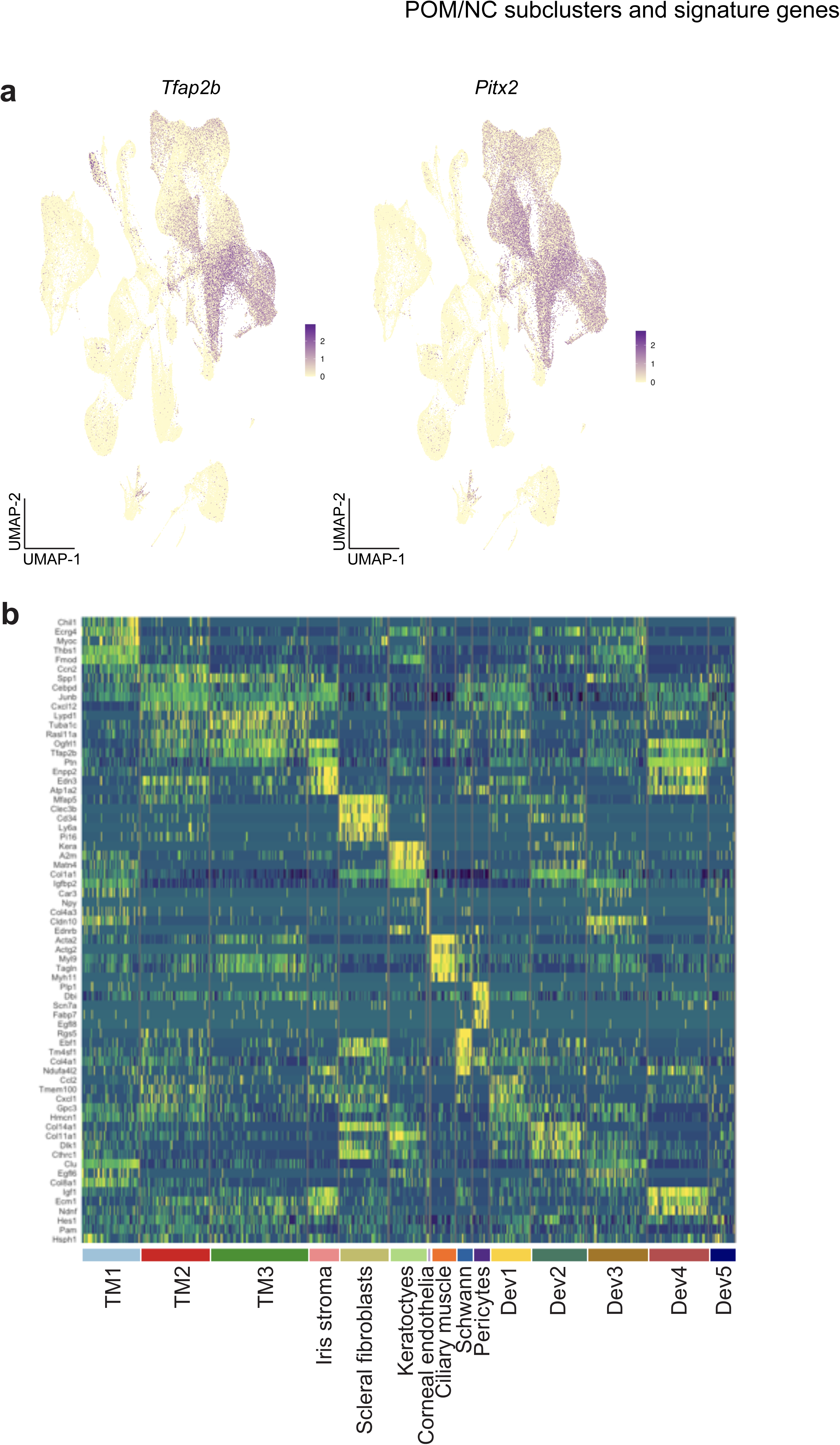
Identification of POM/NC derived cell types. **a)** Visualization of selected marker genes -*Tfap2b* and *Pitx2* - expression plotted on UMAP embedding of all cells. **b)** Heatmap of signature gene expression across single cells arranged by POM-derived cell types.

**Figure S3.**
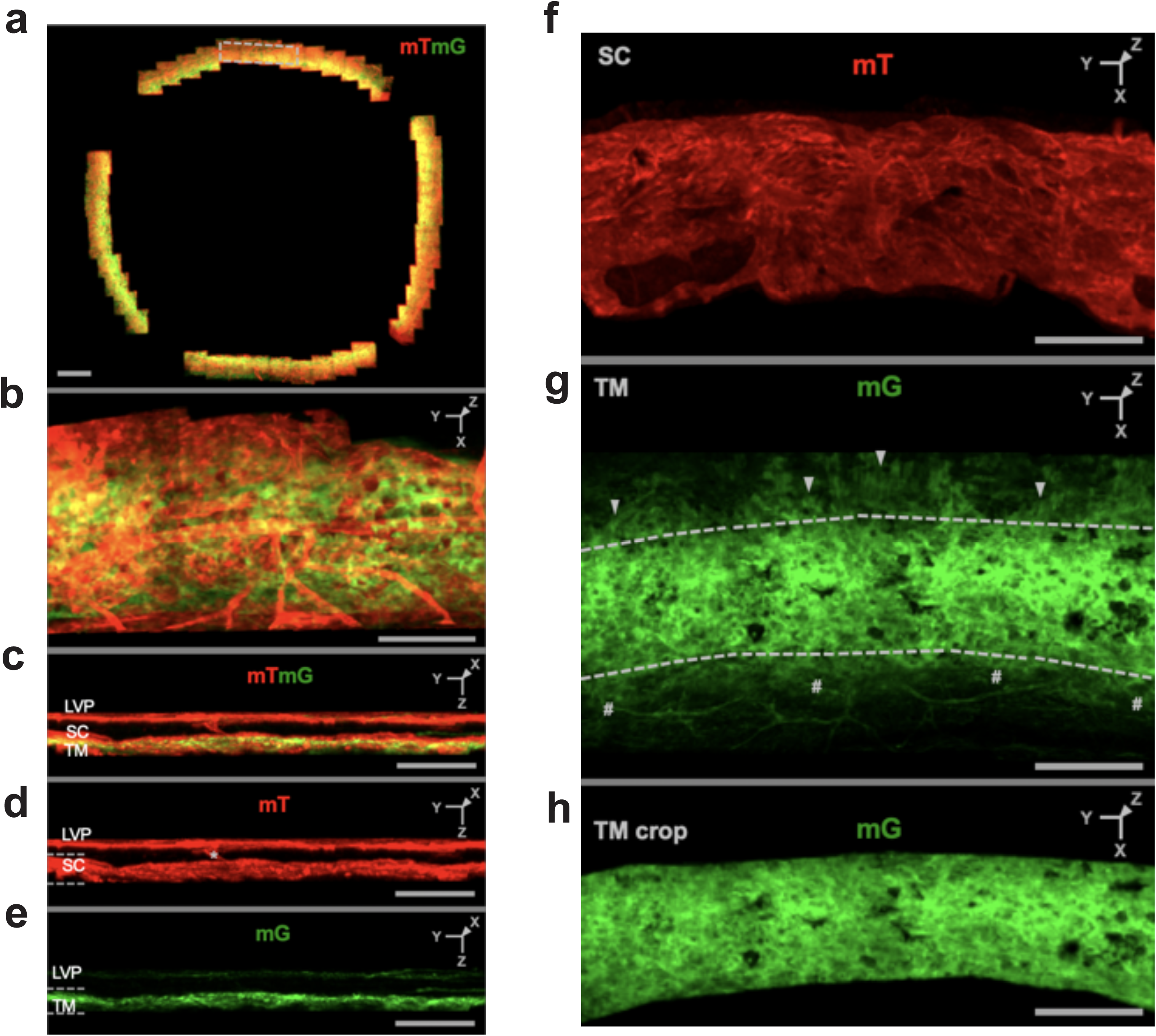
3-D analysis of TM development. This figure describes the anatomic landmarks used to define the TM for Figures 3 and Figures S4 **a)** *En face* view of the entire circumference of the aqueous humor drainage structures and surrounding tissues. Fluorescent labeling was achieved by breeding together the *Wnt1-Cre2* and *mTmG* alleles. This gene combination results in the SC and surrounding vasculature expressing tdTomato (red), while the TM, corneal endothelium, ciliary muscle, and lymphatics express GFP (green; see Methods). The ocular tissue was prepared as a whole mount and imaged through the Z-depth using confocal microscopy. Scale bar = 500 µm**. b)** *En face* view of a cropped region of the AH drainage structures, as indicated by grey lines in (a). The orientation icon in the upper right corner corresponds to that used in subsequent figures**. c-e)** YZ views spanning the full depth of the tissue, from the outer to inner surface of the eye. The limbal vascular plexus (LVP), containing both blood and lymphatic vessels, is located closer to the outer surface of the eye compared to the SC and TM. The LVP was optically removed to view the SC and TM. Collector channels (asterisks) are cropped to within 10 µm of their connection to the outer SC surface. **f-g)** *En face* views of the region between the white dotted lines in (d) and (e), with other regions cropped out. The SC is visible in (f), while the TM and adjacent structures are shown in (g). The TM exhibits the highest GFP signal intensity. At the posterior boundary of the TM, ciliary muscle fibers (arrows) run orthogonally to the TM. At the anterior boundary of the TM, larger, circular cell borders of the corneal endothelium are visible (hashtags). Dotted lines mark the anterior and posterior boundaries of the TM. h) For analysis, the structures surrounding the TM were further cropped out at the dotted lines in (g). All scale bars (b-h) = 100 µm

**Figure S4.**
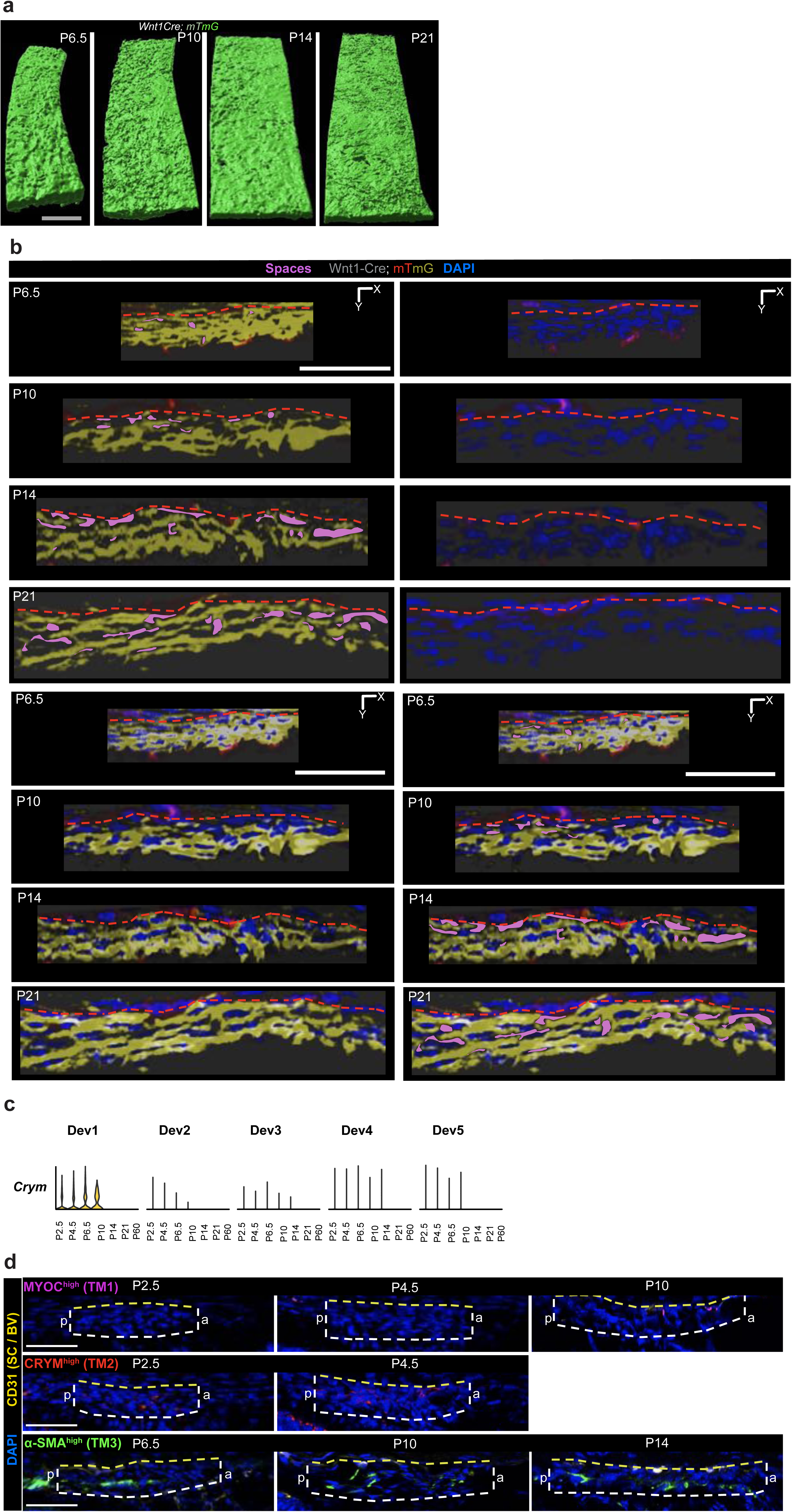
TM cell subtype classification. **a)** View of the AH-facing side of the TM segments shown in Figure 3 (Imaris Surface mode). As development progresses, the structure becomes increasingly organized along its longitudinal axis. **b)** Separated panels of TM XZ optical sections from Figure 3. Empty spaces that lacked both nuclei, (DAPI, blue) and cells (GFP, yellow Methods) are marked (magenta, pseudo-color), red dotted line indicates TM border. **c)** Violin plot showing *Crym* expression across developmental clusters and ages. *Crym* is expressed in the Dev1 cluster from P2.5 to P10, which may explain the detection of CRYM protein in the developing TM prior to P6.5 (before TM2 cells are robustly identified). **d)** Immunofluorescence visualization of TM cell markers shown in Figure 3 at additional developmental ages. MYOC (TM1^Hi^, magenta) is absent in the TM at early stages but becomes detectable by P10. CRYM (TM2^Hi^, red) is present at all examined ages, though early expression may reflect Dev1 identity (see above). α-SMA (TM3^Hi^, green) is consistently expressed across all stages. CD31 and yellow dotted lines demarcate the presumptive SC boundary. White lines outline the TM cell group. a: anterior, p: posterior.

**Figure S5.**
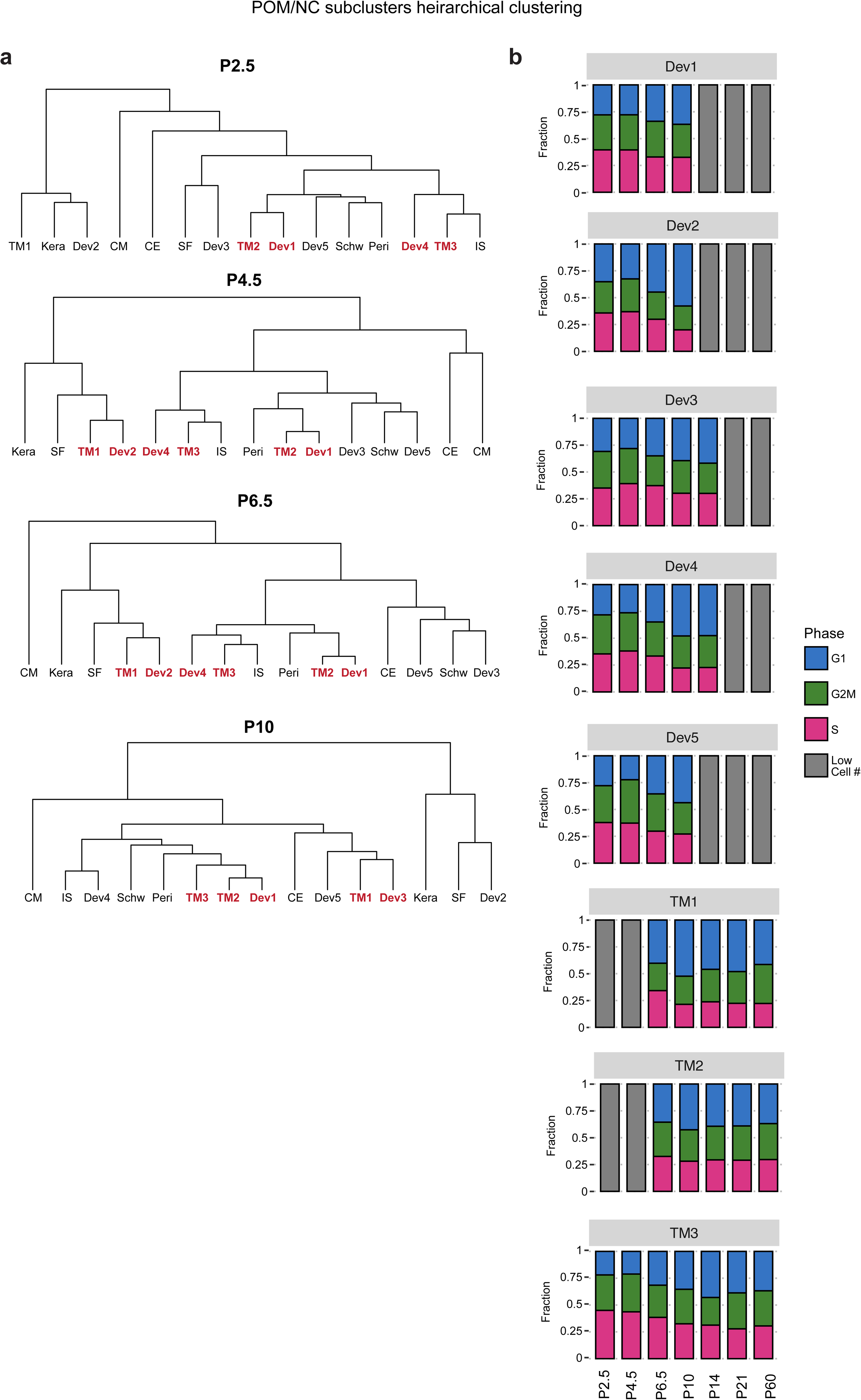
Developmental clusters and TM cell subtypes. **a)** Hierarchical clustering of annotated cell types across P2.5-P10 developmental timepoints for all POM/NC derived cell types. **b)** Barplots demonstrating the proportion of cell cycle phases across sample collection timepoints for the five POM-derived developmental cell types.

**Figure S6.**
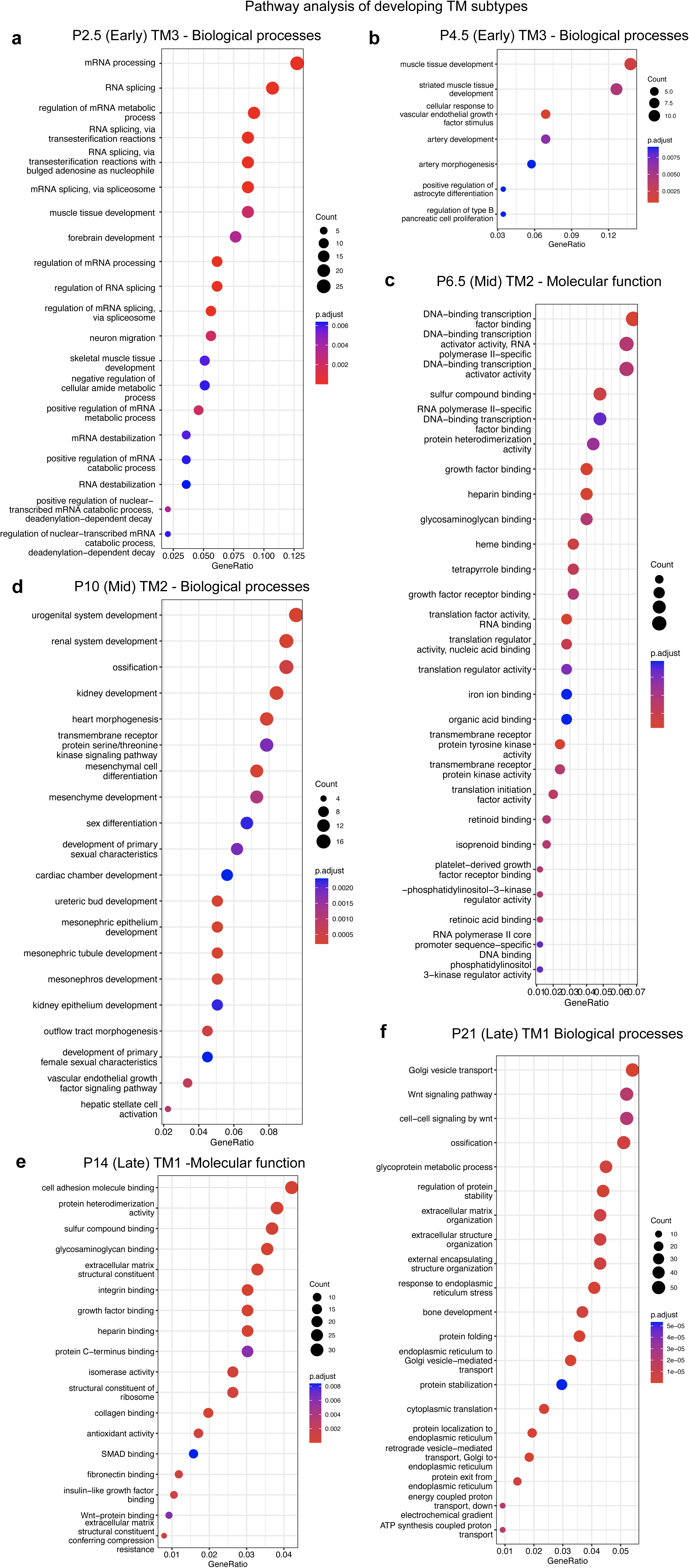
GO enrichment analysis of TM developmental marker genes. **a, b)** GO biological processes of TM3 subcluster marker genes (compared to other POM/NC derived cell types) at P2.5 and P4.5. **c)** GO molecular function of TM2 subcluster marker genes at P6.5. **d)** GO biological processes of TM2 subcluster marker genes at P10. **e)** GO molecular function of TM1 subcluster marker genes at P14. **f)** GO biological processes of TM1 subcluster marker genes at P21.

**Figure S7.**
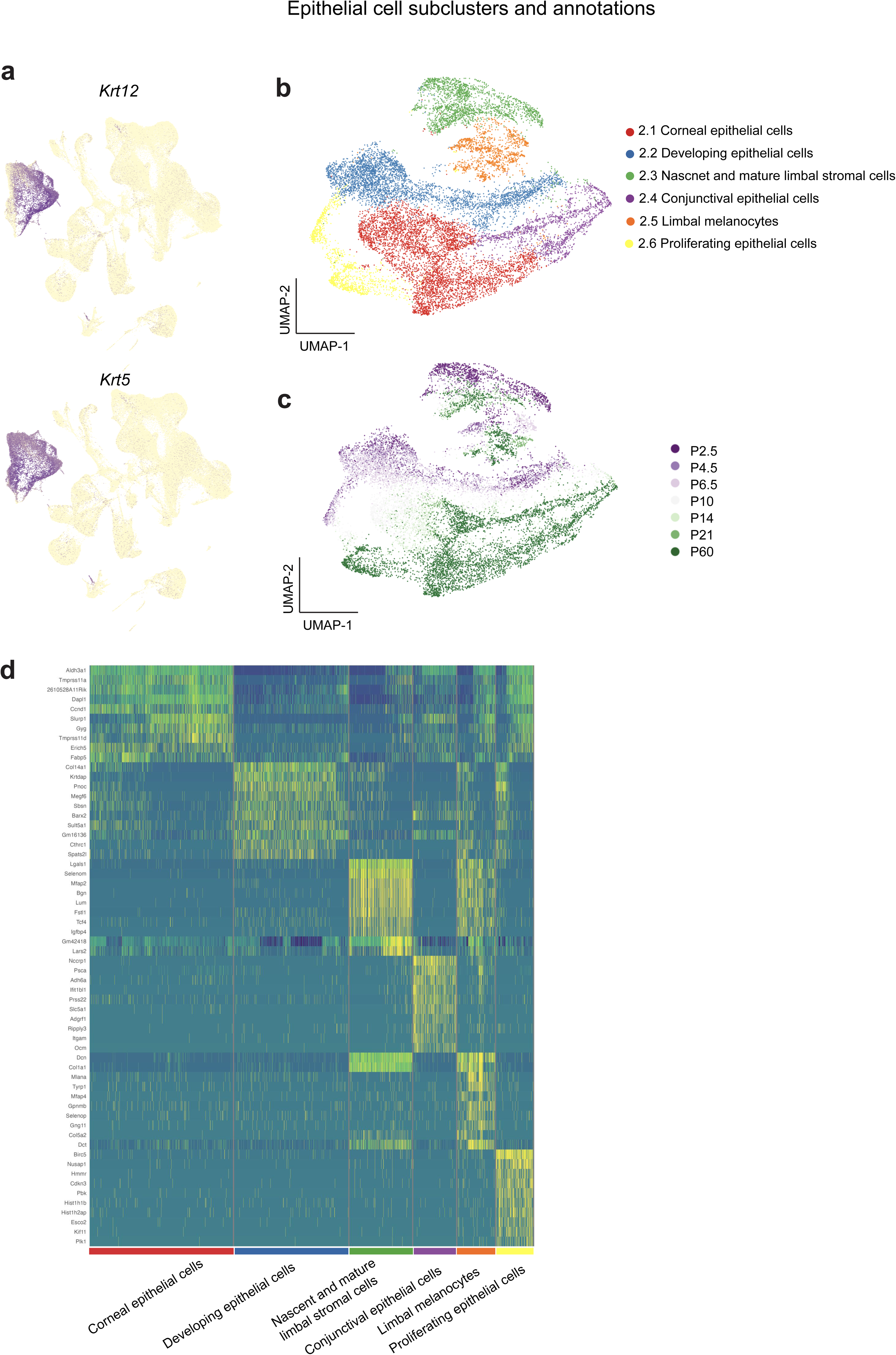
Identification of epithelial cell types. **a)** Visualization of selected marker gene expression plotted on UMAP embedding of all cells. **b)** UMAP representation of epithelial cells colored by subtypes. **c)** UMAP representation of epithelial cells colored by collection timepoint. **d)** Heatmap of signature gene expression across single cells arranged by epithelial cell types.

**Figure S8.**
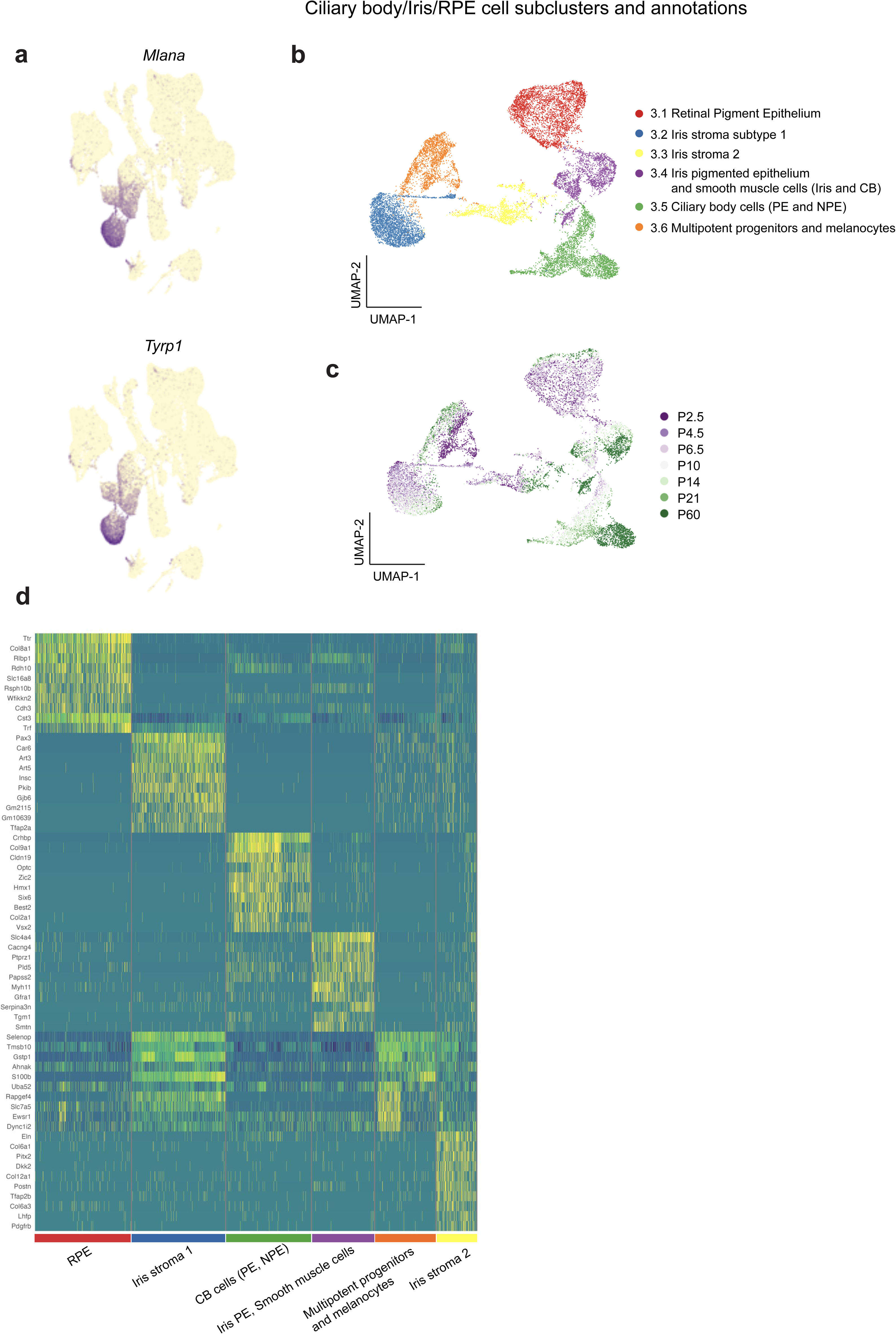
Identification of ciliary body/iris/retinal pigment epithelium (RPE) cell types. **a)** Visualization of selected marker gene expression plotted on UMAP embedding of all cells. **b)** UMAP representation of ciliary body/iris/RPE cells colored by subtypes. **c)** UMAP representation of ciliary body/iris/RPE cells colored by collection timepoint. **d)** Heatmap of signature gene expression across single cells arranged by ciliary body/iris/RPE cell types.

**Figure S9.**
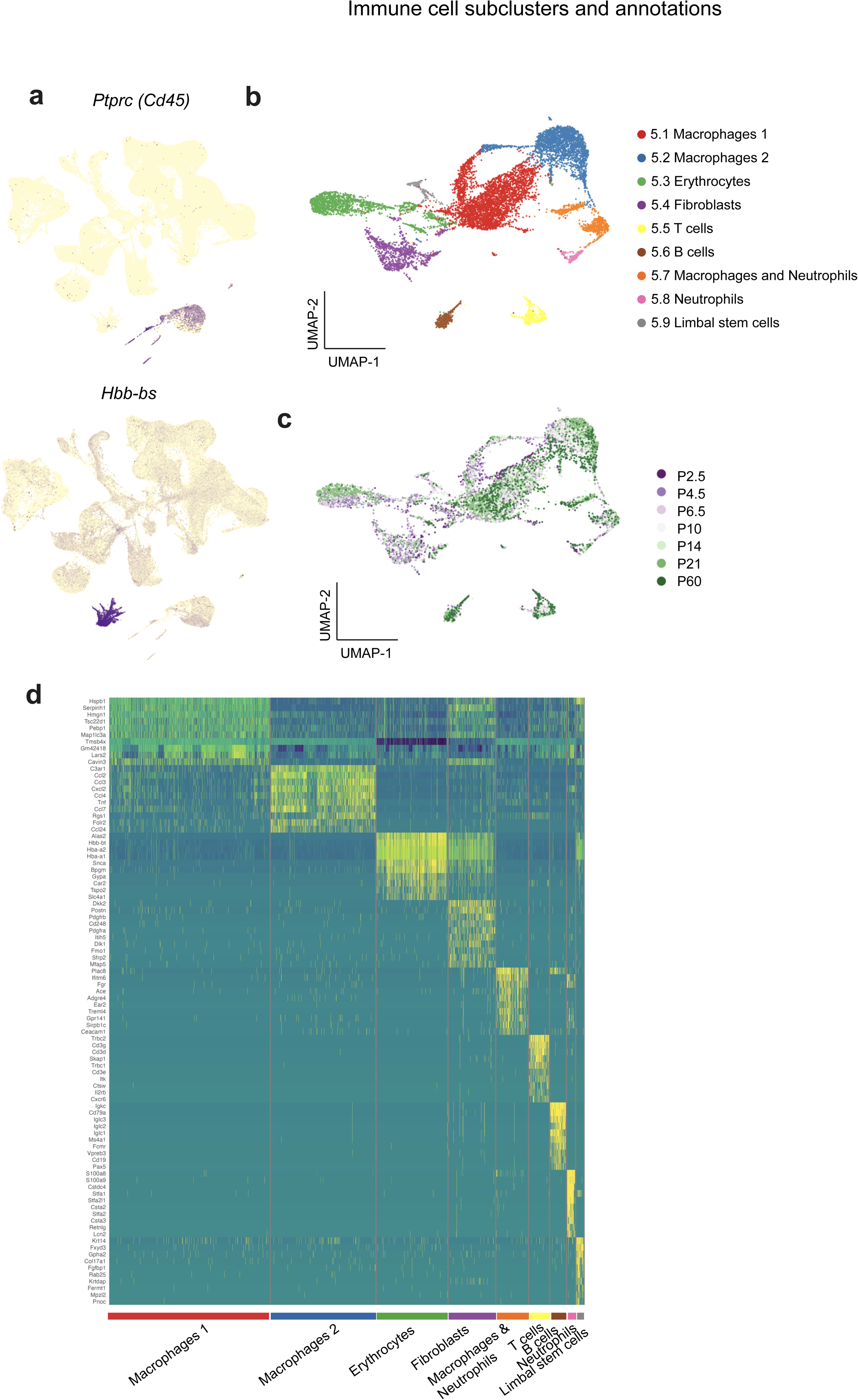
Identification of immune cell types. **a)** Visualization of selected marker gene expression plotted on UMAP embedding of all cells. **b)** UMAP representation of immune cells colored by subtypes. **c)** UMAP representation of immune cells colored by collection timepoint. **d)** Heatmap of signature gene expression across single cells arranged by immune cell types.

**Figure S10.**
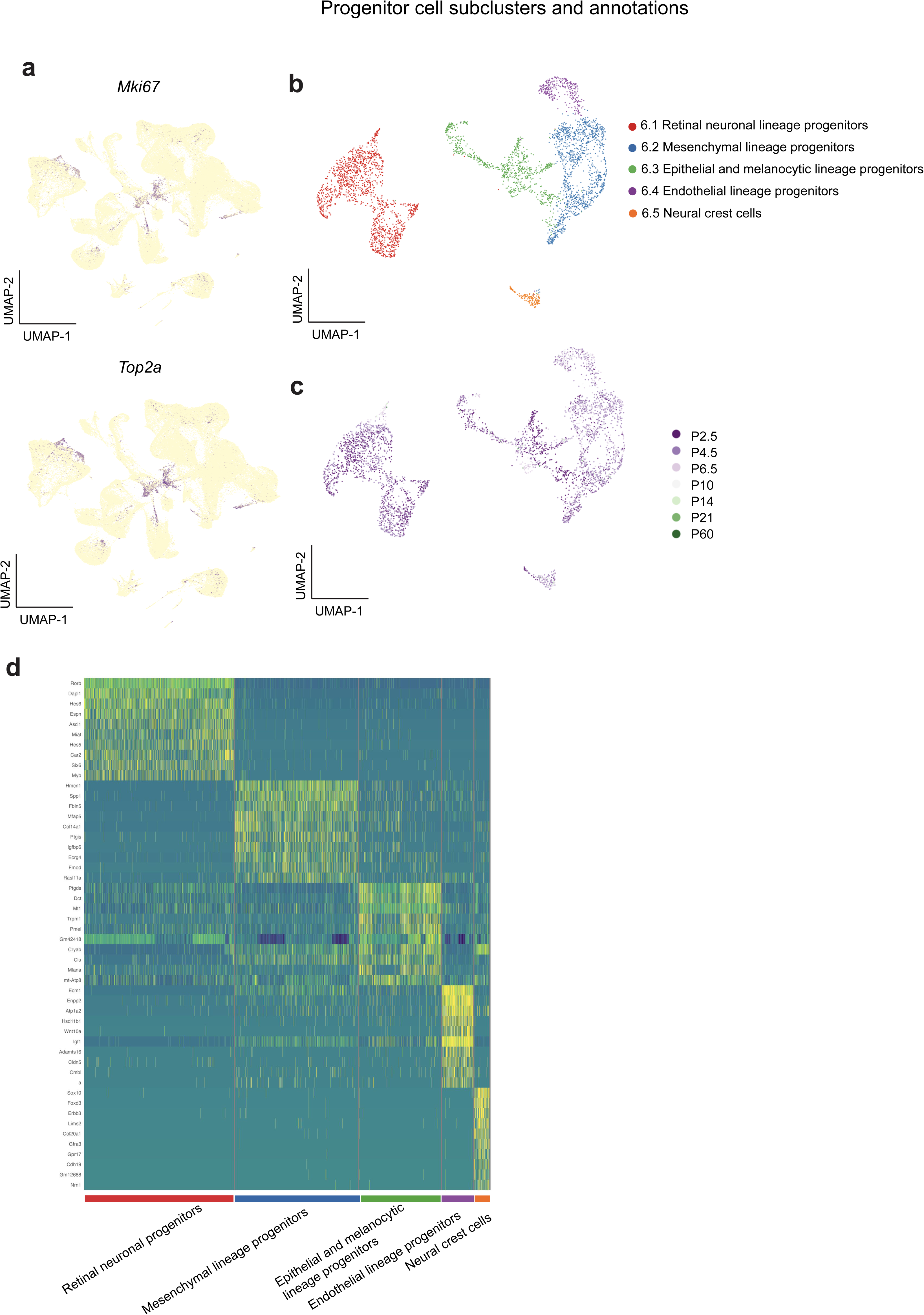
Identification of progenitor cell types. **a)** Visualization of selected marker gene expression plotted on UMAP embedding of all cells. **b)** UMAP representation of progenitor cells colored by subtypes. **c)** UMAP representation of progenitor cells colored by collection timepoint. **d)** Heatmap of signature gene expression across single cells arranged by progenitor cell types.

**Figure S11.**
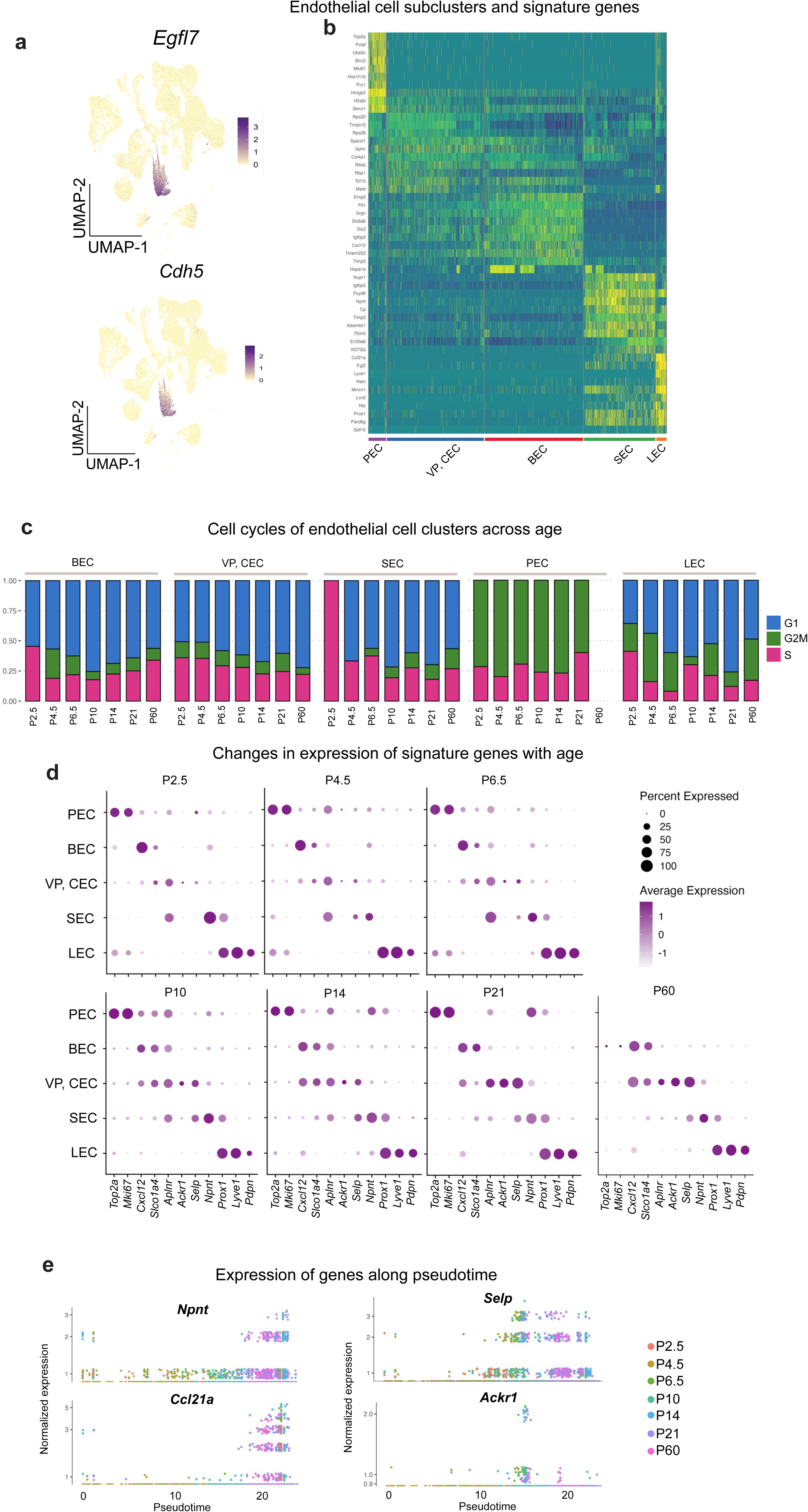
Identification of endothelial cell subtypes. **a)** Visualization of selected marker genes plotted on UMAP embedding of all cells. **b)** Heatmap of signature gene expression across single cells arranged by endothelial cell types. **c)** Barplots demonstrating the fraction of cell cycle phase across sample collection timepoints for the five endothelial cell types. **d)** Dotplots representing the expression of selected marker genes (columns) across sample collection timepoints in each of the five endothelial cell types (rows). **e)** Normalized expression of genes along pseudotime with individual dots colored by chronological time.

**Figure S12.**
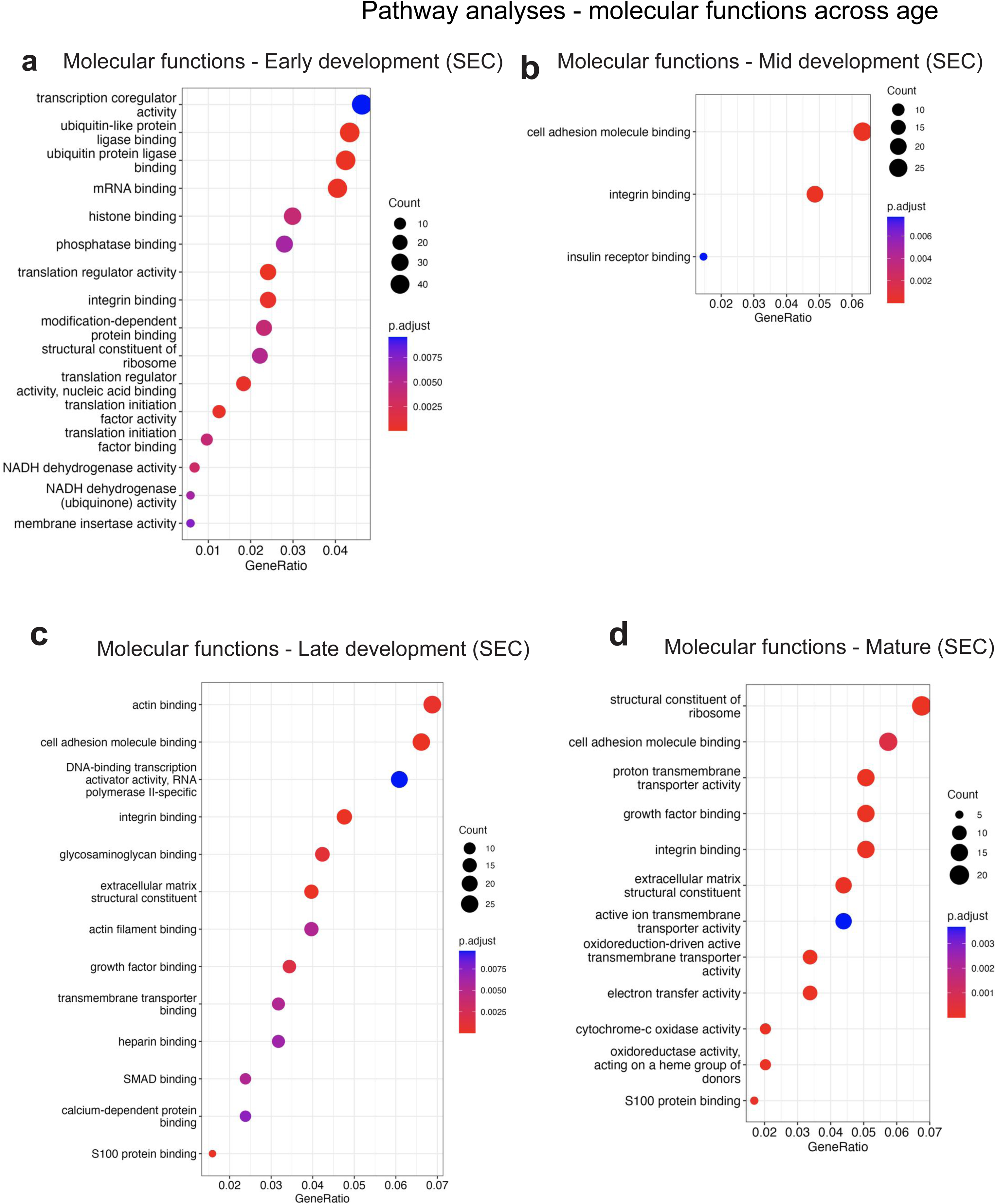
GO molecular pathway enrichment analysis of genes across SEC development. **a-d)** Enrichment of molecular function pathway genes selectively expressed during SEC development.

**Figure S13.**
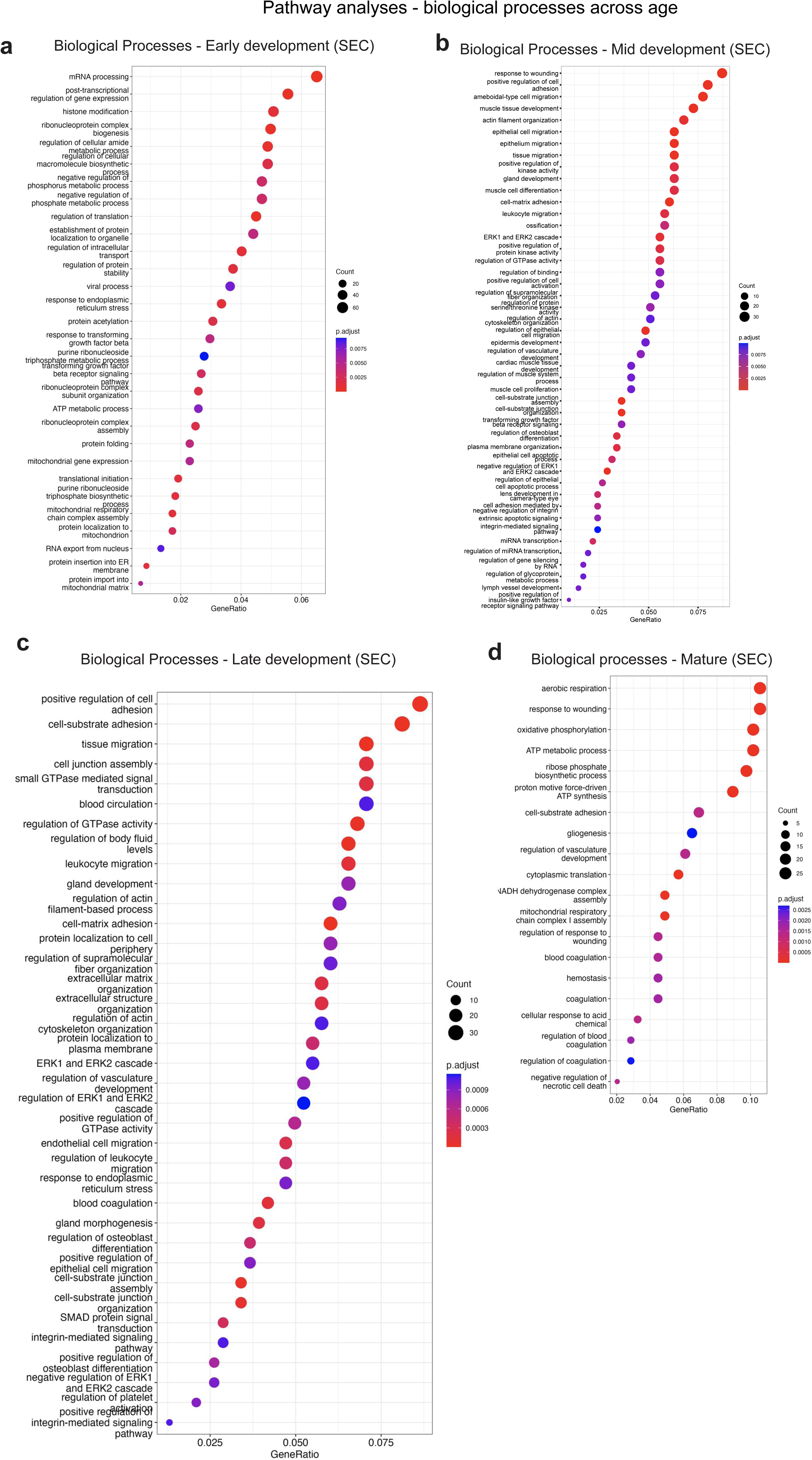
GO biological process enrichment analysis of genes across SEC development. **a-d)** Enrichment of biological process pathway genes selectively expressed during SEC development.

**Figure S14.**
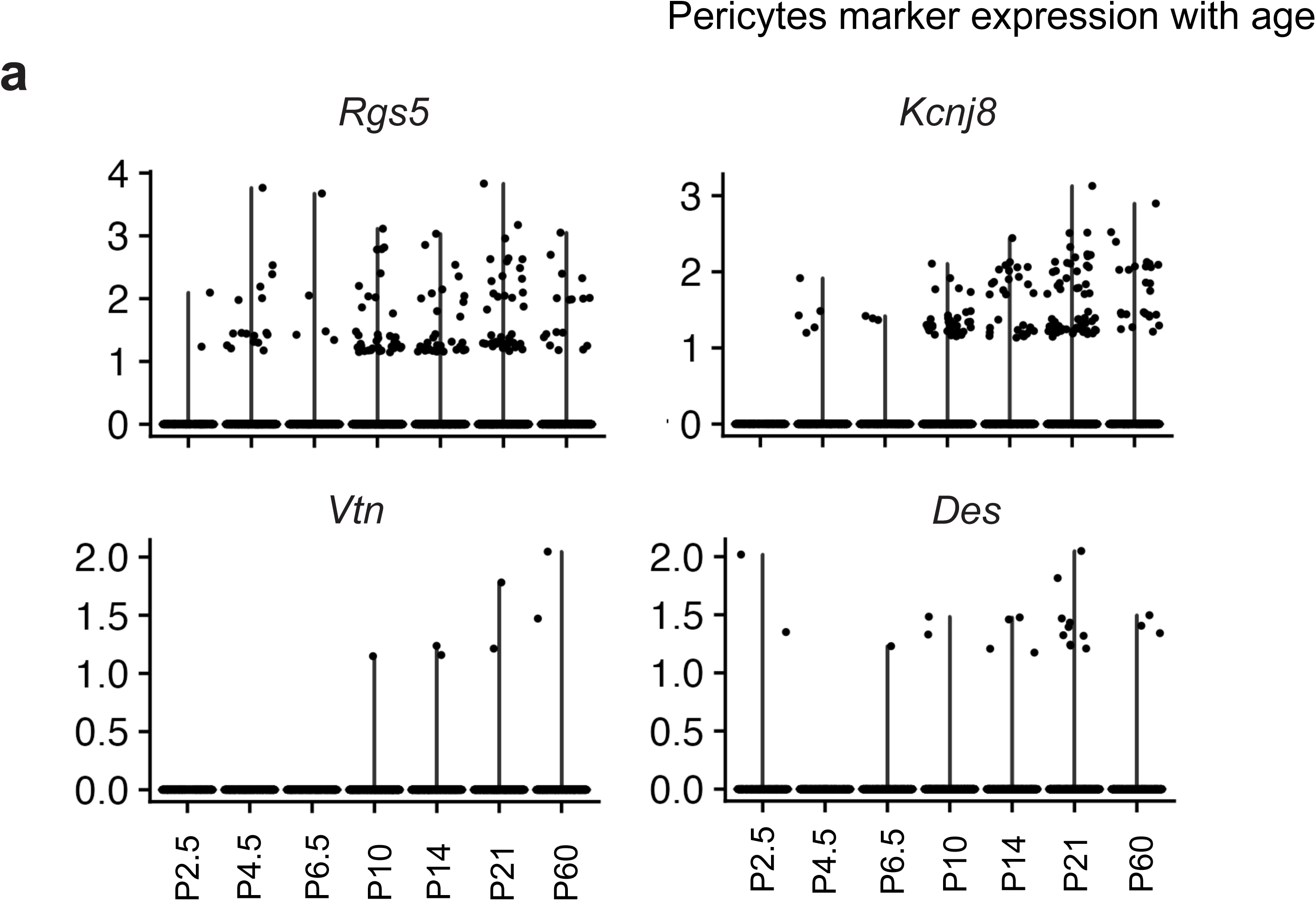
Expression of pericyte marker genes. Violin plots representing the expression profile of four selected pericyte marker genes across sample collection timepoints. Dots represent individual cells.

**Figure S15.**
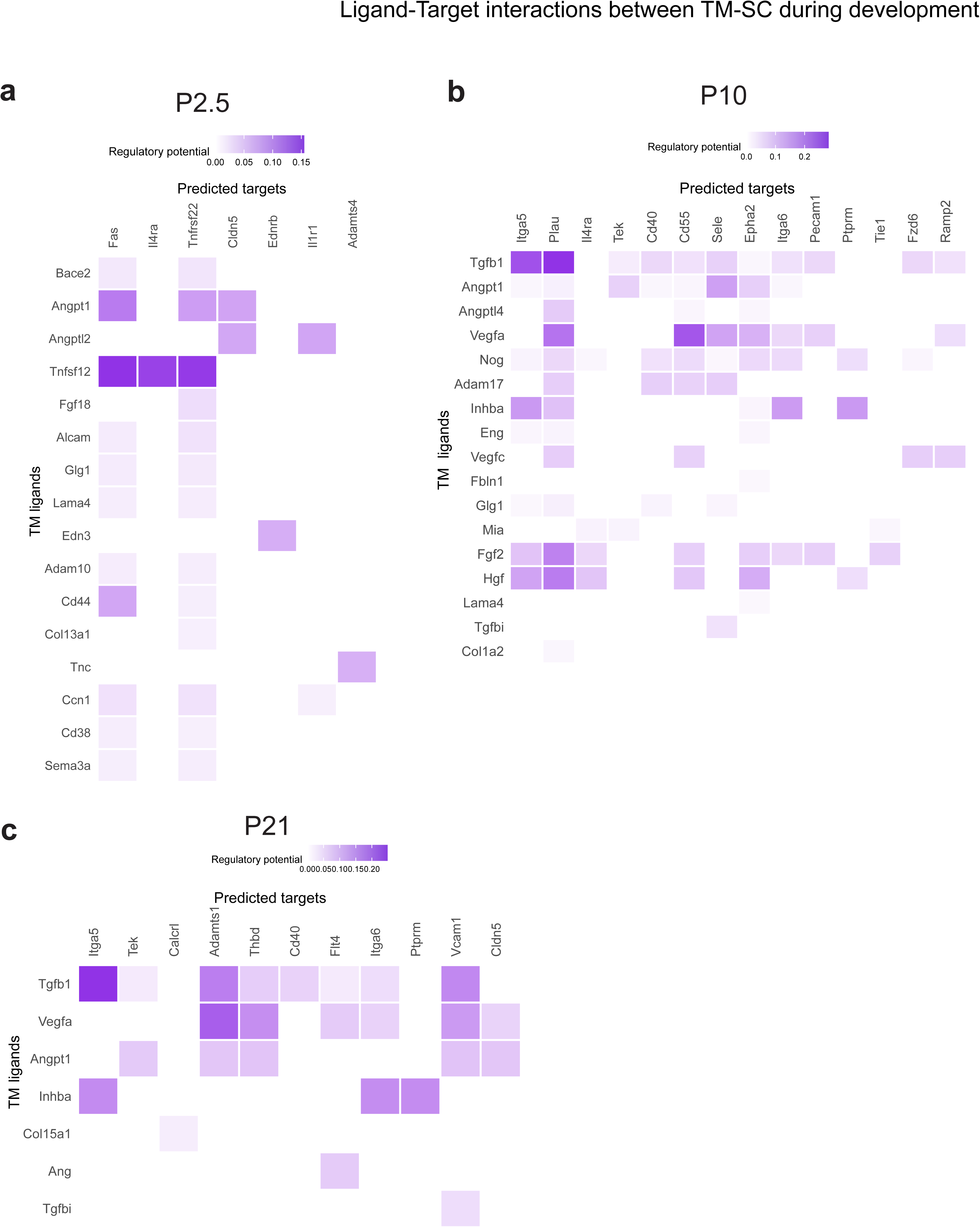
Predicted ligand-target interactions between TM and SC cells during development. Heatmaps represent the regulatory potential between TM ligands (rows) and predicted receptors/targets in SC cells (columns) across three sample collection timepoints (**a:** P2.5; **b:** P10, **c:** P21). NicheNet prioritized extracellular signaling molecules (y-axis) and their predicted downstream direct and indirect targets in the receiving cell population (x-axis), which may include receptors, signaling mediators, ECM components, and transcriptional targets. Color intensity reflects the NicheNet regulatory potential score.

**Figure S16.**
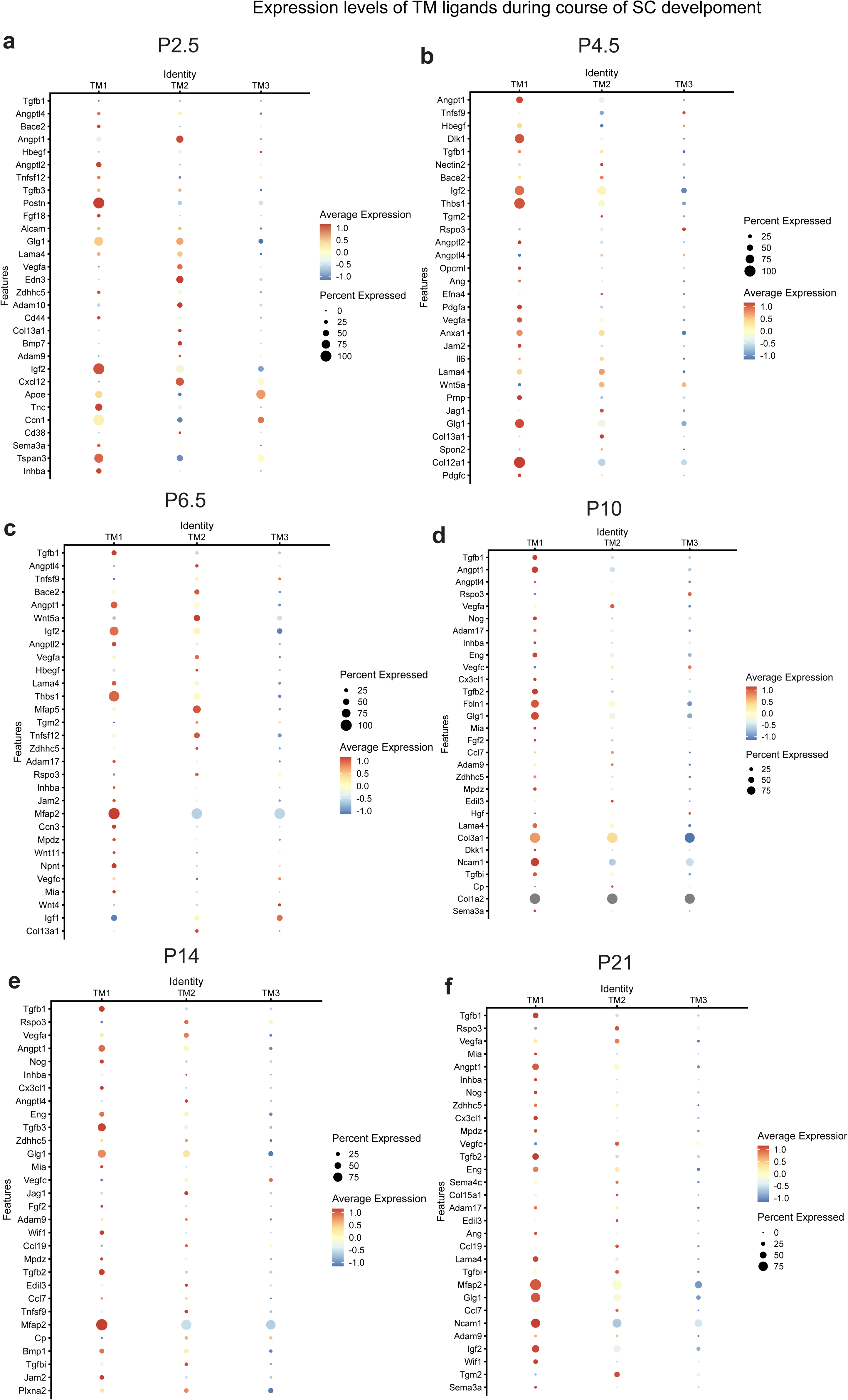
Expression levels of TM ligands during SC development. **a-f)** Dotplots indicating the expression pattern of ligands (ligands) which were predicted to contribute to SC development in each of the three TM cell types (columns) across sample collection timepoints. Dot size is indicative of percent of cells expressing the ligand (not number of cells – due to low numbers of TM1 and TM2 cells at P2.5 and P4.5, the dots should be interpreted with caution at these ages).

**Figure S17.**
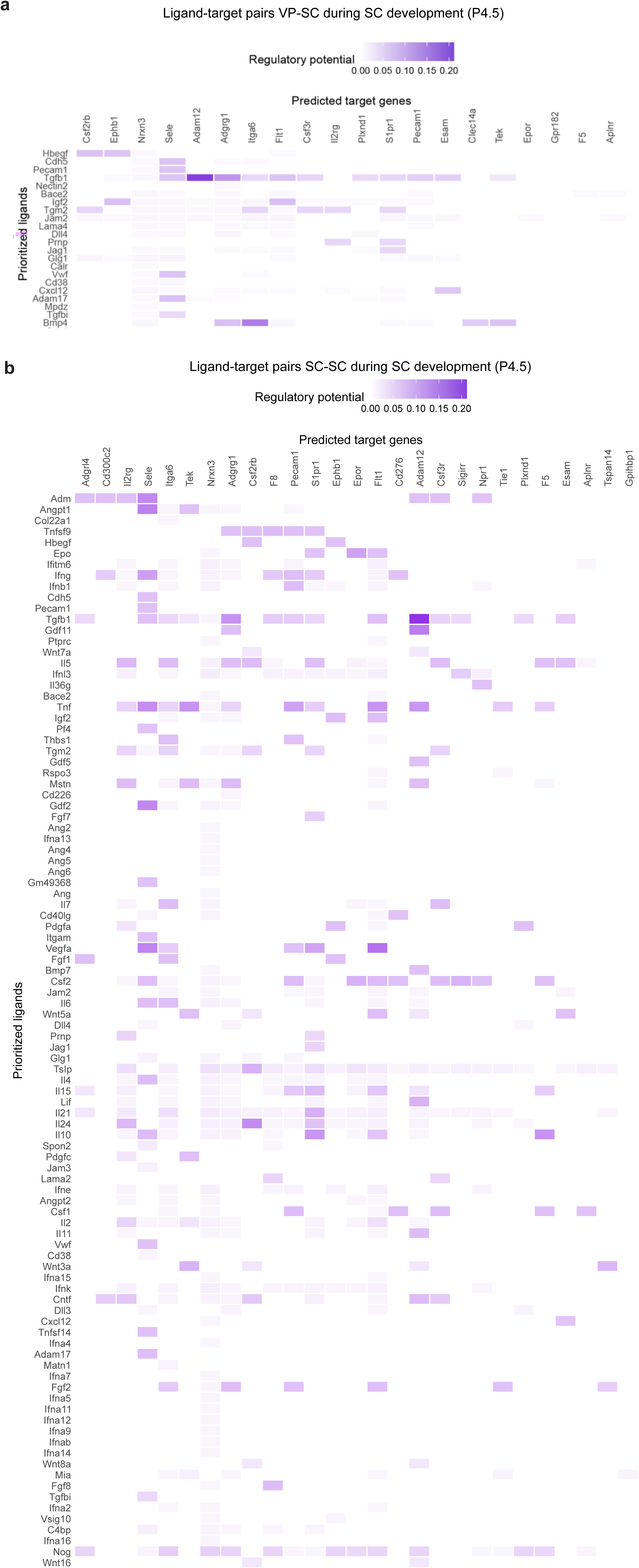
Predicted ligand-target interaction between VP-SC and SC-SC interactions. **a)** Heatmaps representing the regulatory potential between VP ligands (rows) and predicted receptors/targets in SECs (columns) at P4. **b)** Heatmaps representing the regulatory potential between SC ligands (rows) and predicted receptors on SC cells (columns) at P4.

**Figure S18.**
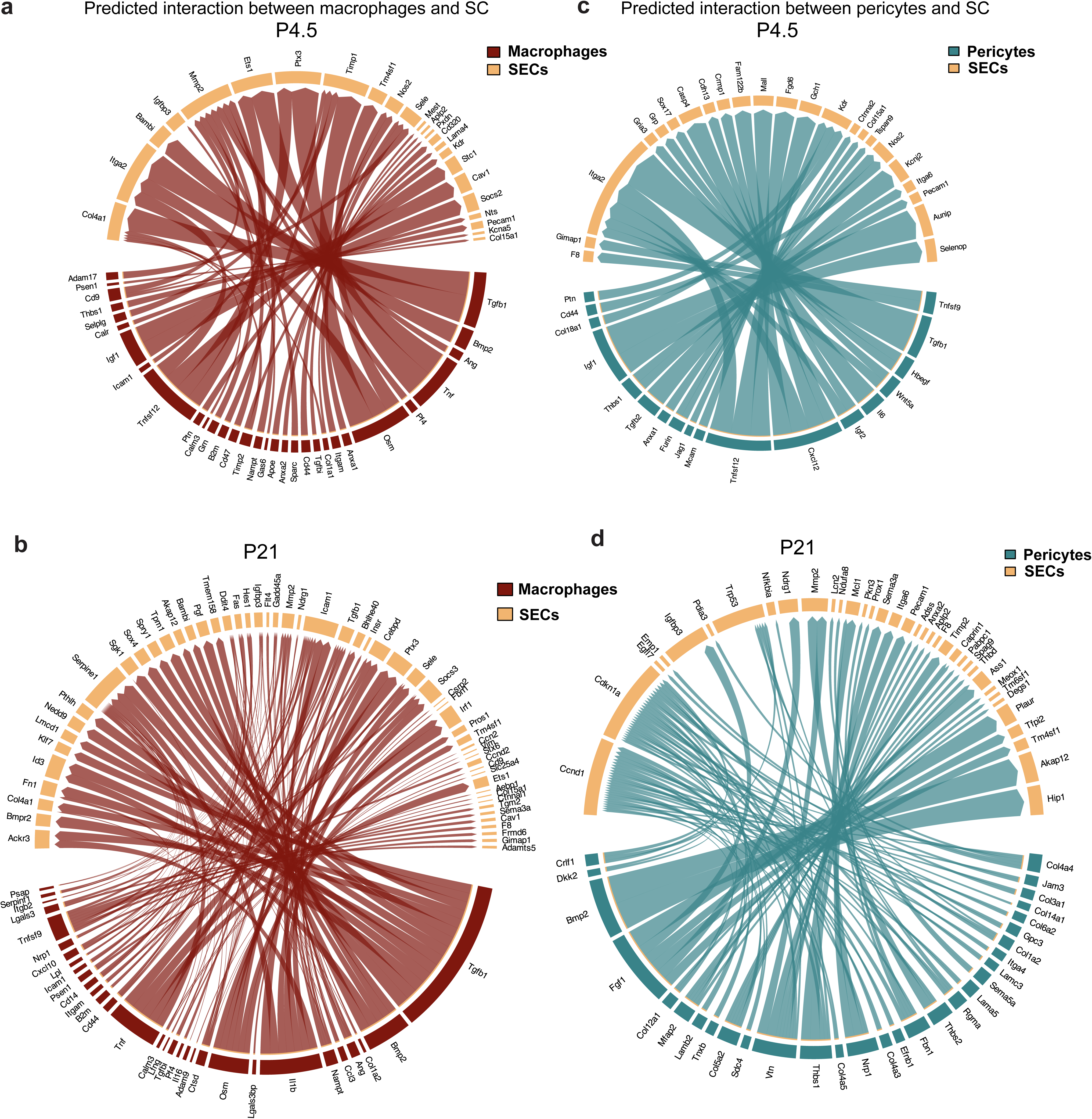
Ligand-target interaction analysis between macrophages/pericytes and SECs. **a, b)** Circos plot demonstrating inferred interactions between macrophages(sender) and SECs (receiver) at P4.5 and P21 **b, c)** Circos plot demonstrating inferred interactions between pericytes (sender) and SECs (receiver) at P4.5 and P21.

**Figure S19.**
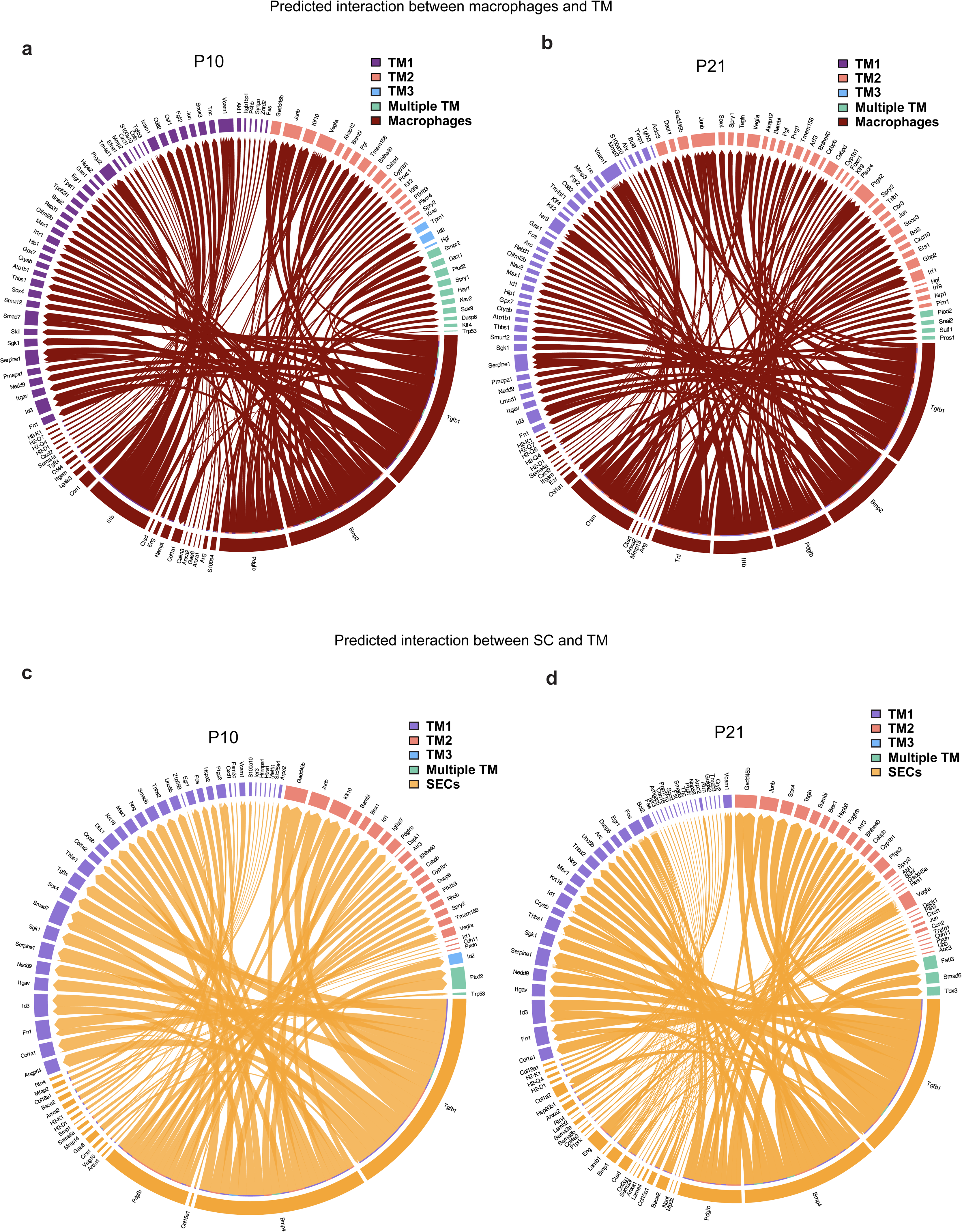
Ligand-target interaction analysis macrophages-TM and SC-TM. **a, b)** Circos plot demonstrating inferred interactions between macrophages(sender) and TM (receiver) cells at P10 and P21 **c, d)** Circos plot demonstrating inferred interactions between SECs (sender) and TM (receiver) cells at P10 and P21.

**Figure S20.**
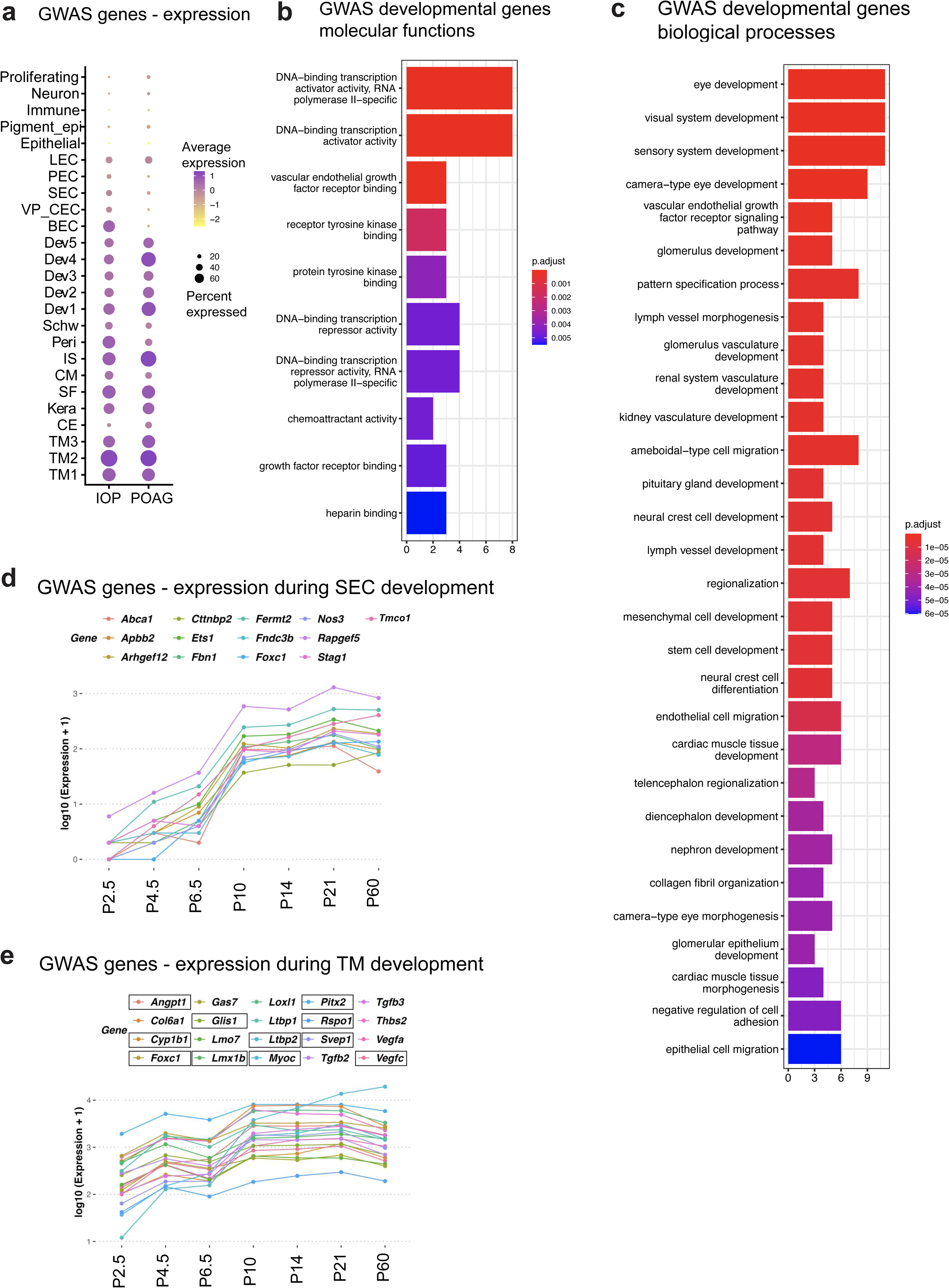
Analysis of expression profile of glaucoma risk genes in SC and TM development. **a)** Dot plot summarizing expression of glaucoma risk genes from IOP and POAG GWAS across all AS cell types. **b, c)** GO molecular function and biological process gene enrichment of genes that cause developmental glaucoma. **d, e)** Average expression level of glaucoma risk genes in SECs and TM cells across sample collection timepoints, developmental glaucoma genes are marked by a rectangular box.

## Supporting information

**Table S1.**
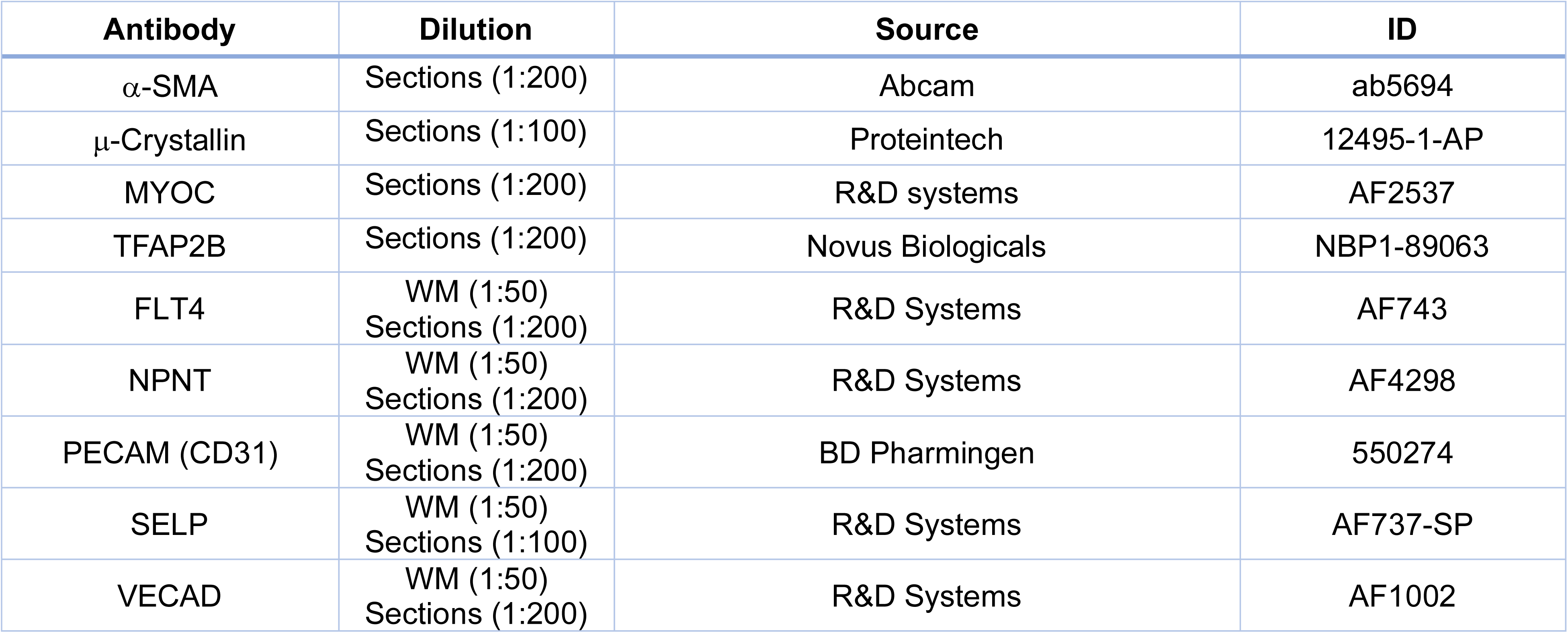
List of antibodies used in the study.

**Table S2.** List of other resources and reagents.

| Resource | Source | ID |
| --- | --- | --- |
| **Animals** |  |  |
| C57BL/6J mice | Jackson Labs | IMSR_JAX:000664 |
| B6.Cg-*E2f1*<sup>Tg(Wnt1-cre)2Sor</sup>/J | Jackson Labs | Stock 022501 |
| B6.129(Cg)-*Gt(ROSA)26Sor*<sup>tm4(ACTB-tdTomato,-EGFP)Luo</sup>/J | Jackson Labs | Stock 007676 |
| **Reagents/Chemicals/Other** |  |  |
| 4% PFA | Thermo Fisher Scientific | Cat no. 50-980-495 |
| 40 and 100 µm Falcon cell strainers | Thermo Fisher Scientific | Cat no. 08-771-1 / 08-771-19 |
| Bovine Serum Albumin | Thermo Fisher Scientific | Cat no. AM2616 |
| Collagenase Type 4 | Worthington Biochemical | Cat no. LS004188 |
| DAPI | Thermo Fisher Scientific | Cat no. 62248 |
| Dulbecco's Modified Eagle Medium (DMEM) | Thermo Fisher Scientific | Cat no. 11965-118 |
| Dulbecco's Phosphate Buffered Saline (PBS) | Sigma Aldrich | Cat no. D8662 |
| Earle's balanced salt solution (EBSS) | Thermo Fisher Scientific | Cat no. 24010-043 |
| Papain Dissociation System | Worthington Biochemical | Cat no. LK003153 |
| Propidium iodide | Thermo Fisher Scientific | Cat no. P1304MP |
| SYTOX green Nucleic Acid Stain | Thermo Fisher Scientific | Cat no. S7020 |

**Table S3:** Markers for developing and adult TM and SC cell types.

| **Markers** | **Target cell type** |
| --- | --- |
| MYOC, CHIL1 | TM1 |
| CRYM, INMT,<br>EDN3 | TM2 |
| α-SMA, TFAP2B,<br>LYPD1, LMX1B | TM3 |
| CXCL1, CCL2,<br>HMCN1 | Dev1 |
| COL14A1, DLK1 | Dev2 |
| CLU, COL8A1 | Dev3 |
| IGF1, ECM1<br>LMX1B | Dev4 |
| HES1 | Dev5 |
| APLNR,<br>CLEC14A | Vascular progenitors |
| IGF1, MMP14 | Early/developing Schlemm's canal cells |
| NPNT, CCL21A,<br>ITGA9 | Schlemm's canal endothelial cells (Inner wall) |
| SELP, LCN2 | Schlemm's canal endothelial cells (Outer wall) |

\*Marker genes are not completely specific to each cell type indicated above. Combination of markers and their levels of expression must be considered. Markers listed above are most highly expressed in the indicated cell types.

## Notes

https://singlecell.broadinstitute.org/single_cell/study/SCP3301/single-cell-characterization-of-anterior-segment-development-cell-types-pathways-and-signals-driving-formation-of-the-trabecular-meshwork-and-schlemms-canal

https://www.ncbi.nlm.nih.gov/geo/query/acc.cgi?acc=GSE315712

### Competing Interest Statement

The authors have declared no competing interest.

### Summary of Updates

Figure 3 has been updated to include quantification of trabecular meshwork volume and space with developing age.

